# Low frequency traveling waves in the human cortex coordinate neural activity across spatial scales

**DOI:** 10.1101/2020.03.04.977173

**Authors:** Vishnu Sreekumar, John H. Wittig, Julio I. Chapeton, Sara K. Inati, Kareem A. Zaghloul

## Abstract

Traveling waves of oscillatory activity are thought to influence neuronal spiking, providing a spatiotemporal frame-work for neural communication. However, no direct link has been established between traveling waves and neuronal spiking in humans. We examined traveling waves in the human lateral temporal lobe by using recordings from intracranial electrodes implanted in twenty participants for seizure monitoring as they performed a paired-associates verbal memory task. We observed ubiquitous low frequency traveling waves across the temporal lobe. While wave occurrence in a broad low frequency range did not differ between successful and unsuccessful memory conditions, in a subset of participants with microelectrode recordings, we found that macro-scale waves co-occurred with micro-scale waves, which in turn were temporally locked to single unit spiking. This temporal coordination between traveling waves at different spatial scales and between waves and neuronal spiking in the human brain suggests a role for traveling waves in neural communication.

## Introduction

Brains are complex dynamical systems that enable us to sense, perceive, and generate meanings in a changing world. Both spontaneous and stimulus-driven brain activity show signatures of flexible transitions between different metastable states (Roberts et al., 2019). One such important state is organized as traveling plane waves of neural activity at both mesoscopic (local, *∼* 1 millimeter, henceforth referred to as “micro-scale”) and macroscopic scales (global, extending over several centimeters, henceforth referred to as “macro-scale”) (Muller et al., 2018). Traveling waves in the brain have been reported extensively in several species, including rabbit, cat, dog, monkey, and turtle (Ermentrout and Kleinfeld, 2001), and have been shown to be functionally relevant, especially in the domains of vision and motor behavior (Rubino et al., 2006; Zanos et al., 2015). Evidence for macroscopic traveling waves in the human brain has primarily come from electroencephalography (EEG) recordings but confounds due to volume conduction have made it difficult to interpret the results (Muller et al., 2018). More recently, intracranial EEG (iEEG) recordings from human clinical patients have revealed macroscopic traveling waves of low frequency oscillatory activity (Muller et al., 2016; Zhang and Jacobs, 2015; Zhang et al., 2018).

Information processing within this complex multi-scale system is thought to rely on coordinated interactions between neuronal populations both within small brain regions and across larger spatial scales. Phase synchronization between participating groups of neurons has been proposed to be the specific mechanism underlying such interactions (Varela et al., 2001). Traditionally, the focus of such analyses has been on a precise temporal alignment of oscillatory activity across brain regions, but based on recent studies, it appears that phase offsets organized systematically in space are a common feature of both cortical and subcortical neural activity. These propagating spatiotemporal patterns include traveling plane waves (Lubenov and Siapas, 2009; Zhang et al., 2018), spiral waves (Muller et al., 2016), and other complex waves (Rubino et al., 2006; Townsend et al., 2015). A cortical repertoire of such time-lagged phase synchrony patterns may provide a more flexible mechanism for functional interactions between neural populations, and recent evidence has demonstrated that these patterns can transiently appear over durations of as little as tens to hundreds of milliseconds. For example, visual information processing induces “perceptual echoes” in the form of posterior-anterior alpha traveling waves that recur periodically across cycles once every *∼* 100 ms (Lozano-Soldevilla and VanRullen, 2019). Similarly, spontaneous cortical activity measured using resting-state MEG transiently organizes into phase-coupled networks, and these transient states are visited repeatedly with the dwell times in each state lasting an average of 50 to 100 ms (Vidaurre et al., 2018). While there are different types of propagating wave patterns as described above, we focus on propagating plane waves in the current study. These are waves defined by wave fronts that are approximately linear constant phase contours.

In addition to providing a framework for time-lagged phase synchronization, traveling waves may also play an active role in information processing by influencing the probability of neuronal firing. In this manner, traveling waves could coordinate spiking activity between brain regions in space and time (Wu et al., 2008). Studies have suggested such a link between traveling waves and neuronal activity at smaller scales (micro-scale and single unit levels) by using high frequency activity at the macro-scale as a proxy for spiking (Bahramisharif et al., 2013). In addition, previous evidence has explicitly demonstrated human single unit spiking activity locked to the phase of theta/alpha and delta oscillations (Chapeton et al., 2019; Jacobs et al., 2007; Rutishauser et al., 2010), raising the possibility that traveling waves within these frequencies may coordinate spiking activity. However, there is currently no direct evidence that traveling waves coordinate spiking activity in humans. The presence of traveling waves that coordinate local spiking activity, if identified, would be consistent with theoretical accounts of communication between brain regions. Both empirical evidence and computational models suggest that communication may rely on coherent low frequency oscillations that coordinate higher frequency bouts of spiking activity in different areas of the brain (Chapeton et al., 2019; Fries, 2005; Haegens et al., 2011). Traveling waves propagating across brain regions could provide a mechanistic account for how such communication and coordination of neuronal activity could arise.

Here, we investigated how traveling waves are orchestrated across spatial scales in the human brain by examining concurrent iEEG and microelectrode (MEA) recordings from the human temporal cortex. We detected traveling plane waves at both spatial scales using optical flow, which estimates motion (instantaneous velocities) from sequences of images. In this case, images were instantaneous phase maps of neural activity in a low frequency band that was chosen by first computing the frequencies within individuals at which maximum coherence was observed between electrodes. We examined traveling waves within a relatively broad range of low frequencies that enabled us to investigate the precise temporal relation between traveling waves across spatial scales. At the macro-scale, we found traveling waves with speeds and directions consistent with previous reports (Zhang et al., 2018) and with durations that lasted anywhere between tens of milliseconds to a few hundred milliseconds depending on the extent of cortical space covered by the waves. We also identified micro-scale traveling waves in the MEA recordings captured from the human middle temporal gyrus, and obtained direct evidence that low frequency micro-scale traveling waves and single unit spiking activity are tightly coupled. Macro-scale wave times were correlated with both micro-scale wave times and spike times. Taken together, these results establish a link between macro-scale traveling waves, micro-scale traveling waves, and single-unit spiking activity in the human brain, suggesting that traveling waves may play a role in information processing in the brain by coordinating spiking activity both across different brain regions and within local patches of the cortex.

## Results

### Macro-scale traveling waves

We investigated macro-scale cortical traveling waves in twenty participants with drug-resistant epilepsy who under-went surgery for placement of intracranial electrodes for seizure monitoring (intracranial EEG, iEEG; Fig. 1A). We examined traveling waves while participants were attentive and repeatedly engaged in the same behavior in order to explore the potential role of traveling waves in human cognition. We therefore restricted our analyses to data that had been collected while participants performed a verbal paired-associates memory task (see Methods) that required similar cognitive processing during every trial of the task rather than analyze data from random times during the day (Fig. 1B). Participants studied 377 ± 46 word-pairs across experimental sessions and correctly recalled words cued by the other word in a given pair in 0.31 ± 0.05 proportion of the trials with a reaction time of 1968 ± 113 ms. In order to examine traveling waves, we first computed spectral coherence between every electrode pair on the temporal macro-electrode grid in each participant to identify the frequencies most likely involved in coordinating activity across larger-scale brain regions. We found that the average coherence spectrum over all pairs exhibited a prominent peak in most participants (see Methods). This peak coherence frequency was generally in the high theta/low alpha range (range = 2-14 Hz; mean across participants = 7.67 ± 0.53 Hz; Fig. 1C). The presence of these peaks indicated that there were consistent phase relationships between oscillations at different locations at these frequencies. We were primarily interested in understanding the precise temporal relation between wave-like activity patterns in these frequencies at multiple spatial scales and single unit spiking activity. Given the time-frequency resolution tradeoffs inherent in filtering iEEG data, however, it was important to use a sufficiently broad spectral filter capable of preserving this temporal precision (see Methods). We therefore used a 4 *−* 14 Hz FIR filter to filter the iEEG data in a single broad low frequency band that captures the peak in coherence across participants (see Methods).

**Figure 1.**
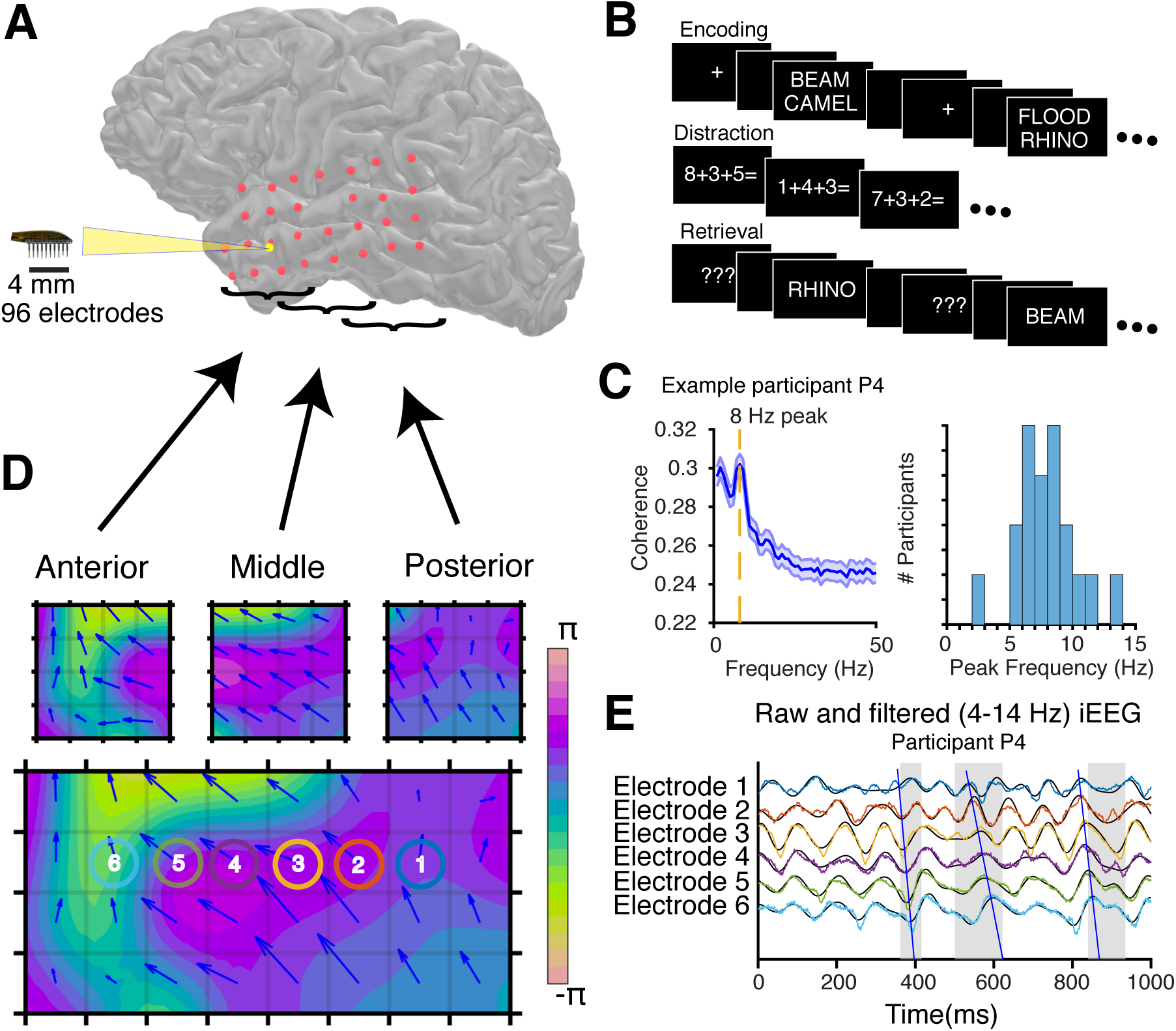
Detecting traveling waves at the macro- and micro-scales through iEEG and microelectrode array (MEA) recordings from the temporal lobe. (*A*) Locations of the 8 *×* 4 iEEG macro-electrode grid (red) and MEA (yellow) on the temporal lobe of an example participant. (*B*) During encoding, pairs of words are sequentially presented on the screen. Subsequently during retrieval, one word from each pair is randomly chosen and presented, and participants are instructed to vocalize the associated word. (*C*) The coherence spectrum of an example participant averaged across trials (the blue shaded region is the SEM) is shown on the left and a histogram of peak coherence frequencies across participants is shown on the right. (*D*) The macro-electrode grid is divided as shown into three overlapping square grids. Traveling waves are detected separately on each square sub-grid. Traveling waves that are concurrently detected on all three sub-grids are combined to identify waves that travel the entire extent of the 8 *×* 4 grid. An example of such a “long-range” traveling wave from participant P4 is shown here (see Supplemental Movie S1 for the corresponding movie). (*E*) iEEG waveforms (filtered traces (4-14 Hz) are shown in black) from six electrodes (indicated in *D*) that lie along the path of the example traveling wave shown in *D*. The second gray highlight denotes the duration of this traveling wave and the blue line tracks the sequentially arriving peaks of the EEG waveform at these six electrodes. Also shown by gray highlights are two other wave epochs detected within the same 1 s period, one before (the blue line tracks the troughs), and one after the example traveling wave.

We then used an algorithm based on optic flow to determine epochs where activity in this low frequency band propagated across the cortical surface as traveling plane waves. To ensure that the algorithm is not biased in any direction due to asymmetry of the electrode arrangement, we limited our analysis only to electrodes arranged in a grid on the cortical surface and applied the detection algorithm separately to three overlapping 4 *×* 4 square subgrids overlying the lateral temporal lobe (Fig. 1D; see Methods). Once waves were detected separately on the anterior, middle, and posterior square subgrids, we identified waves that propagated in the same direction concurrently across the three subgrids, and designated these as long-range waves (*∼* 8 cm) that traveled across the full length of the 8 *×* 4 temporal lobe grid (Fig. 1D; Supplemental Movie S1). To demonstrate the lagged phase relationship that is typically observed during a traveling wave, we picked six electrodes along the direction of an exemplar traveling wave and plotted the iEEG traces of those electrodes (Fig. 1E). Within an epoch identified by the algorithm as a traveling plane wave, there is a clear progressive phase lag between adjacent electrodes.

We confirmed that the detection algorithm only identifies traveling waves when there are true lagged phase relationships between multiple channels on the grid by comparing the detected traveling waves to those detected using surrogate iEEG data. We generated phase-randomized surrogate iEEG data with the same autocorrelations as in the true data but with reduced cross-correlations between electrode channels (reduced by approximately 25%; Supplementary Fig. S1A,B; see Methods). We only partially distorted the cross-correlations in order to provide a more conservative test. The true data exhibited a significantly greater number of traveling waves per second detected on each subgrid than the surrogate data across participants (11.74 ± 0.10 versus 9.91 ± 0.23; *t*(19) = 8.93, *p* = 3.15 *×* 10*^−^*^8^; Supplementary Fig. S1C). We found that the average estimate of instantaneous power across all electrodes in each subgrid was significantly greater during wave epochs compared to non-wave epochs (*t*(19) = 21.92*, p* = 6 *×* 10*^−^*^15^; see Supplementary Fig. S1D) which demonstrates the observed waves were not affected by poorly measured phase estimates when power is low. This difference in power is consistent with previous proposals that oscillations propagate as traveling waves (Zhang et al., 2018).

After removing IEDs and other artifacts (also see Supplementary Information for an analysis of the relationship between wave occurrence and epilepsy), we found that traveling waves across participants were more frequent on the anterior subgrid compared to the posterior subgrid (10.29 ± 0.02 versus 10.03 ± 0.02 waves per second; *t*(19) = 2.59, *p* = 0.018). We also found that traveling waves detected on the anterior subgrid lasted slightly but consistently longer than those on the posterior subgrid 28.0 ± 1 ms versus 26 ± 1 ms; *t*(19) = 3.46, *p* = 0.003).

We examined the direction of the identified traveling waves and found that they generally propagated from a posterior to anterior direction in any given local area of the temporal cortex (Rayleigh’s test *p <* 0.01 for all participants and both anterior and posterior subgrids; Figs. 2A,B). Across participants, the distribution of wave directions was consistently and significantly non-uniform (length of mean resultant vector of all anterior temporal lobe (ATL) wave directions *r* = 0.20 ± 0.03; and of posterior temporal lobe (PTL) wave directions *r* = 0.12 ± 0.01; Hodges-Ajne test for non-uniformity within each participant and local area of the temporal lobe, max *p* = 9 *×* 10*^−^*^8^). Traveling waves in the anterior subgrid were more directional than those in the posterior subgrid (*t*(19) = 3.69, *p* = 0.002).

**Figure 2.**
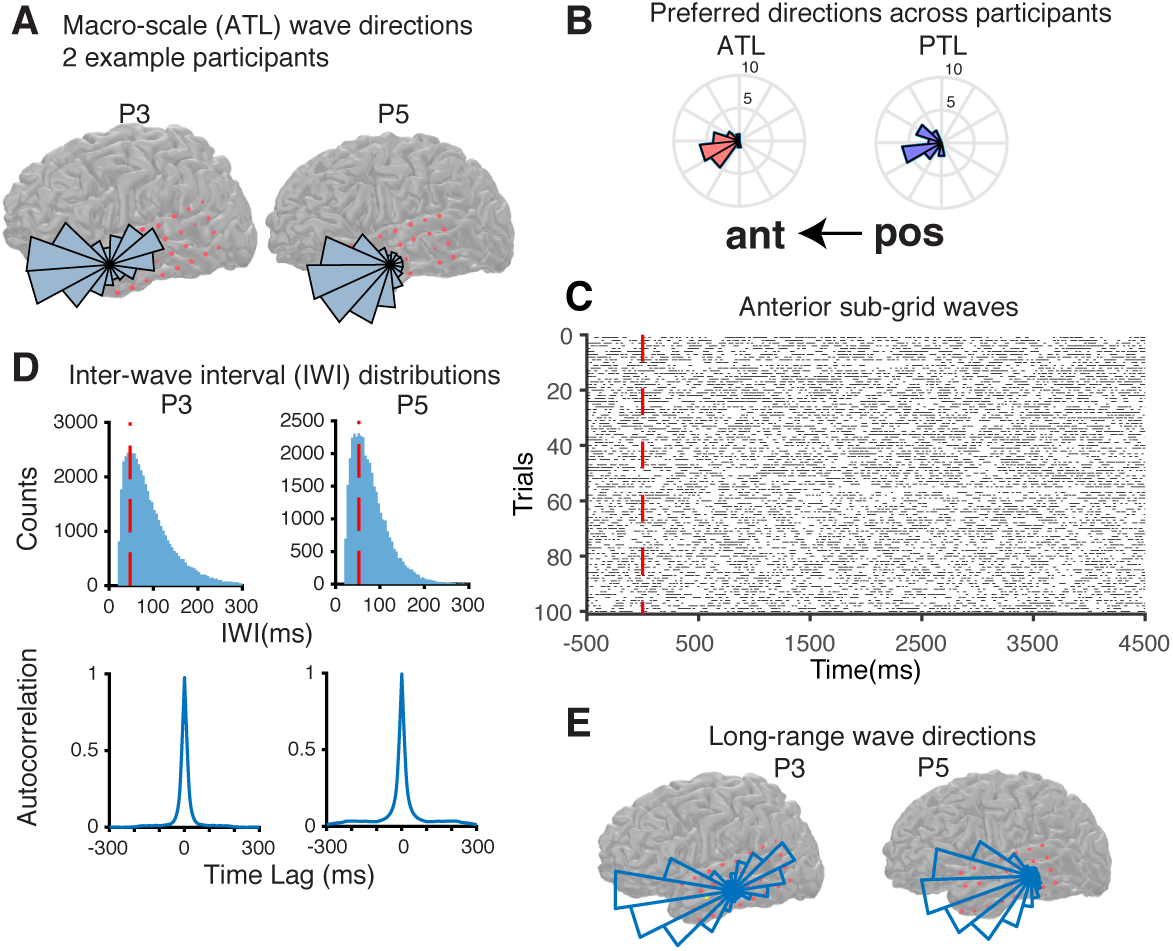
Macro-scale traveling waves on macro-electrode subgrids. (*A*) The distribution of directions of macro-scale waves on the anterior temporal lobe in two example participants. (*B*) The preferred directions of traveling waves in the anterior temporal lobe (ATL) and posterior temporal lobe (PTL) across all 20 participants showing a posterior-anterior preference. *(C)* A raster demonstrating the ubiquity of waves detected on the anterior square sub-grid in an example participant. Each of 100 trials is represented in each row, and the time points during which a traveling wave is detected on the sub-grid are indicated by horizontal lines. The red vertical dotted line indicates the time at which the word-pair appeared on the screen. *(D)* Inter-wave interval distributions for waves in two representative participants and the corresponding autocorrelation in wave times. (*E*) Distribution of directions of long-range cortical traveling waves in two example participants showing a strong preference for the posterior-anterior direction.

Given prior reports of periodic traveling waves (Lozano-Soldevilla and VanRullen, 2019), we investigated whether macro-scale traveling waves were periodic. In each participant, we identified traveling waves during both encoding and retrieval trials of the experimental session. On each temporal lobe 4 *×* 4 subgrid, we found ubiquitous traveling waves that did not appear either periodic or time locked to the onset of the word pairs (Fig. 2C, 100 encoding trials from an example participant. We then examined the inter-wave intervals (IWIs) measured as the time between the beginning of one wave to the beginning of the next. In all participants, we found a similarly shaped distribution of IWIs for traveling waves (Fig. 2D). The shapes of these IWI distributions closely resemble the IWI distributions that would arise assuming a Poisson process with a refractory period that is equal to the wave durations in our data (see Supplementary Fig. S2A). Fig. 2D also includes the autocorrelation in wave times showing no signatures of periodicity. Together, these data suggest that traveling waves in a broad 4 *−* 14 Hz low-frequency range in the human cortex are ubiquitous, are not periodic, and do not appear to be modulated by the onset of the word pairs during encoding or the onset of the cue word during retrieval.

While macro-scale traveling waves detected on the iEEG subgrids were common, less frequently we observed waves that propagated over the full length of the temporal macro-scale electrode grid. These macro-scale long-range traveling waves preferred the posterior-to-anterior direction over anterior-to-posterior in all but one participant (two example participants shown in Fig. 2E; Rayleigh’s test of uniformity *p <* 7.96 *×* 10*^−^*^12^ and *z*-statistic *>* 25.32 across the 19 participants who showed this posterior-anterior preference). Across all participants, 72.0 ± 2.8% of the long-range waves traveled in the posterior-anterior direction (*t*(19) = 4.32*, p* = 0.00037, paired *t*-test). Long-range posterior-anterior waves occurred at the rate of 0.34 ± 0.03 per second whereas the corresponding rate for anterior-posterior waves was 0.12 ± 0.01 per second. Long-range traveling waves that propagated in the posterior-anterior direction were significantly longer lasting than those traveling in the anterior-posterior direction (*t*(19) = 5.33, *p* = 3.9 *×* 10*^−^*^5^; 75 ± 2 ms posterior-anterior waves versus 66 ± 2 ms anterior-posterior waves; Supplementary Fig. S2B shows distributions of macro-scale long-range wave durations in six example participants). Long-range waves propagated at 0.55 ± 0.03 m/s (see Methods). We did not find a significant relation between these long-range 4 *−* 14 Hz traveling waves and successful associative memory formation (see Supplementary Information).

### Micro-scale traveling waves

We wanted to understand how the identified macro-scale traveling waves relate to spatiotemporal patterns of neural activity at a smaller spatial scale. In six participants, we had the opportunity to examine micro-scale traveling waves in the local field potentials captured by a grid of 10 *×* 10 micro-electrodes (microelectrode array, MEA; see Methods). Similar to the macro-scale, we computed the spectral coherence using the microelectrode data and found that five out of six participants had coherence peaks at the micro-scale that were within 1 Hz of the coherence peaks calculated at the macro-scale. The remaining participant had a difference of *∼* 2 Hz. These frequencies were all within the 4 *−* 14 Hz range and we therefore used the same 4 *−* 14 Hz FIR filter to filter the microelectrode data. Because each MEA is a symmetric grid of micro-electrodes, we directly applied the optic flow detection algorithm to data recorded on each MEA in order to identify micro-scale traveling waves without the concern that geometric asymmetry would bias our results (see Methods).

An example micro-scale wave traveling from an inferior-posterior to superior-anterior direction is shown in Fig. 3A (and Supplemental Movie S2). We found fewer micro-scale traveling waves per second than the macro-scale waves detected using iEEG contacts (1.66 ± 0.62 micro-scale waves per second across participants). The mean duration of micro-scale waves was 30 ± 1 ms (Supplementary Fig. S3A). Though less frequently, we did observe longer lasting waves on the order of 100 ms. The mean maximum duration across subjects was 171 ± 22 ms. We found that micro-scale traveling waves were approximately five times slower than their larger macro-scale counterparts, propagating at 0.11 ± 0.007 m/s. When we examined the IWIs between successive micro-scale waves in each participant, we found that the distributions of IWIs for the micro-scale waves were not uniform (Supplementary Fig. S3B). However, we repeated the simulation procedure described earlier assuming a Poisson random process with a refractory period but now chose a rate parameter that matched the measured rate of micro-scale wave occurrence in each individual and found that the locations of peaks obtained from the simulations were not different from those of the empirically observed peaks (*t*(5) = *−*0.78*, p* = 0.47). While the peak locations were predicted by a random generation process, Supplementary Fig. S3C shows that such a random Poisson process with refractoriness cannot by itself account for the shapes micro-scale IWI distributions like it did at the macro-scale suggesting a more complex generation process at the micro-scale. The autocorrelations in micro-scale wave times shown in Supplementary Fig. S3D also do not show signatures of periodicity.

**Figure 3.**
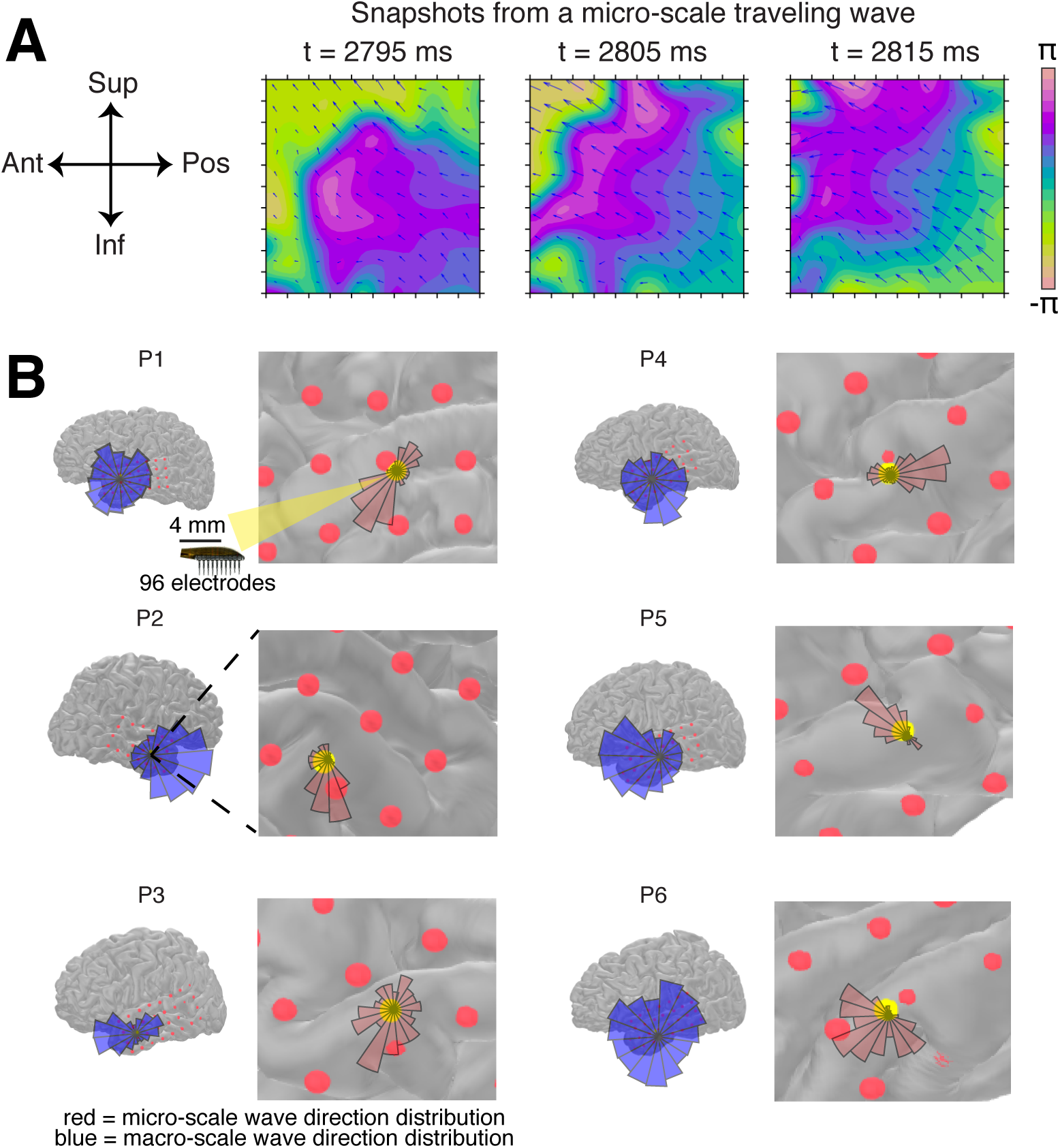
Micro-scale traveling waves. (*A*) An example of a micro-scale traveling wave captured at three different time points on a 96-channel microelectrode array (MEA). Supplemental Movie S2 shows the full wave. (*B*) Distributions of macro-scale wave directions are shown in blue on the brain plot on the left for each participant. A zoomed-in view shows the location of the MEA (yellow dot) and the micro-scale wave direction distribution (red) along with adjacent gyri and sulci for each participant.

Similar to their macro-scale counterparts, the micro-scale waves tended to propagate along specific directions (Fig. 3B). For each participant, we found that the directions of micro-wave traveling waves were significantly non-uniform (Hodge-Ajne omnibus test *p ≤* 6.85 *×* 10*^−^*^39^ for all tests; mean resultant vector length of direction distributions = 0.37 ± 0.06). The directions of propagation of the micro-scale waves, however, were not strictly aligned with the directions of the macro-scale waves (Fig. 3B). Together with the lower rate of occurrence and the slower speed of waves observed at the micro-scale compared to the macro-scale, these data raise the possibility that the waves detected at the micro-scale are not the same waves detected at the macro-scale, and that the directions of these micro-scale waves may be a consequence of the specific architecture and connectivity structure of the local cortical patch (Fig. 3B also shows a magnified view that shows adjacent sulci and gyri).

### Single unit spiking is locked to micro-scale traveling waves

We examined single unit spiking data from the MEA locked to the beginning of each micro-scale traveling wave (Fig. 4A; see Methods), and found a clear peak in spiking activity around wave onset in two participants who had a relatively large number of well isolated single units recorded on the MEA (Fig. 4B; Supplementary Fig. S4A; see Supplementary Table 1 for counts of well isolated units recorded in different participants and experimental sessions). One concern when examining locked spiking data is that spike artifacts and subsequent filtering may give rise to the observed coordination between micro-scale traveling waves and spiking activity. We therefore confirmed that the micro-scale wave-locked spike raster exhibited increased spiking activity following wave onset even after removing spikes from the LFP time series (Supplementary Fig. S4B; see Methods).

**Figure 4.**
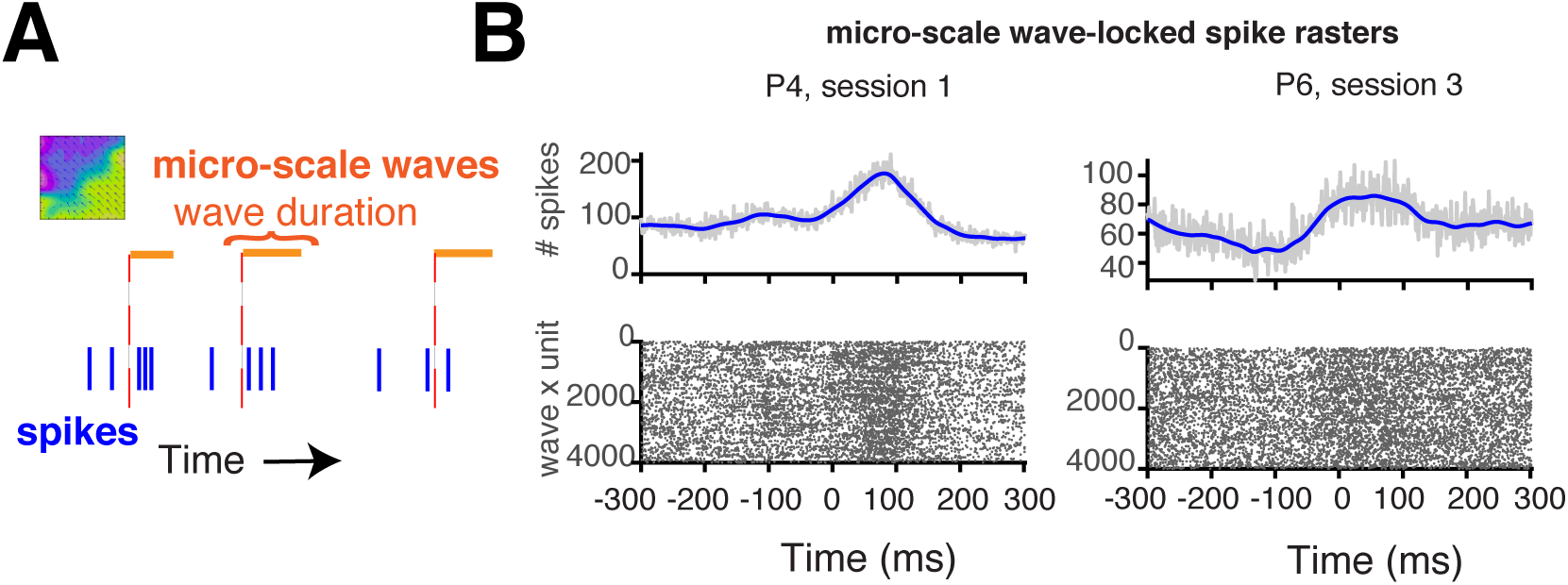
Single unit spiking activity locked to micro-scale traveling waves. (*A*) A schematic of the analysis to identify the relation between micro-scale waves and neuronal spiking. Each orange line represents the duration of a detected micro-scale wave in time. We captured the time points of single unit spiking (blue lines) locked to the beginning of each of these waves in both backward and forward directions. (*B*) Two example participants show increased likelihood of spiking after the onset of micro-scale traveling waves. Spike rasters accumulated over waves and units are shown on the bottom. The sum of spikes at each millisecond is summarized in the plots on top. The blue trace represents a smoothed version (using a Gaussian window of 50 ms) of the continuous time series of summed spiking events.

In order to examine this relation between micro-scale traveling waves and spiking activity across all participants, we computed high frequency broadband (HFB) power (70 *−* 200 Hz, see Methods) as a proxy for multiunit spiking activity (Manning et al., 2009) (see Methods). We confirmed that the spike times identified in this manner using HFB power were a good proxy for local multiunit spiking by computing the cross-correlation between HFB spike events and single unit spike events in every participant. (Supplementary Fig. S5A). In most participants, we found a clear peak in HFB spiking activity that was locked to and followed the onset of micro-scale traveling waves (see Supplementary Fig. S5B). To further test whether micro-scale waves co-occurred with spikes, we computed the cross-correlation between spike times and micro-scale wave times and found a greater positive relationship for the true but not surrogate data (see Methods) in 4 out of 6 participants (corrected *p <* 0.05; Supplementary Fig. S6A). This set of analyses taken together suggests that micro-scale waves and single unit spiking activity are temporally coordinated in a majority of participants.

### Relation between traveling waves at the macro- and micro-scales

Traveling waves detected at the macro and micro scales appear to propagate in different directions and at different speeds, suggesting that they are separate wave phenomena. Yet it is possible that the wave phenomena at the two different spatial scales are temporally related. We examined this possibility directly by computing the cross-correlation between macro-scale and micro-scale wave times averaged across all trials, in each participant (Fig. 5A; see Methods). Two example participants’ data are shown in Fig. 5B. We found a greater peak in the average cross-correlogram between micro-scale and macro-scale wave times at around time = 0 in five out of six participants in the true data compared to surrogate data (corrected *p <* 0.05; Supplementary Fig. S6B), suggesting significant temporal coordination between waves at the different scales.

**Figure 5.**
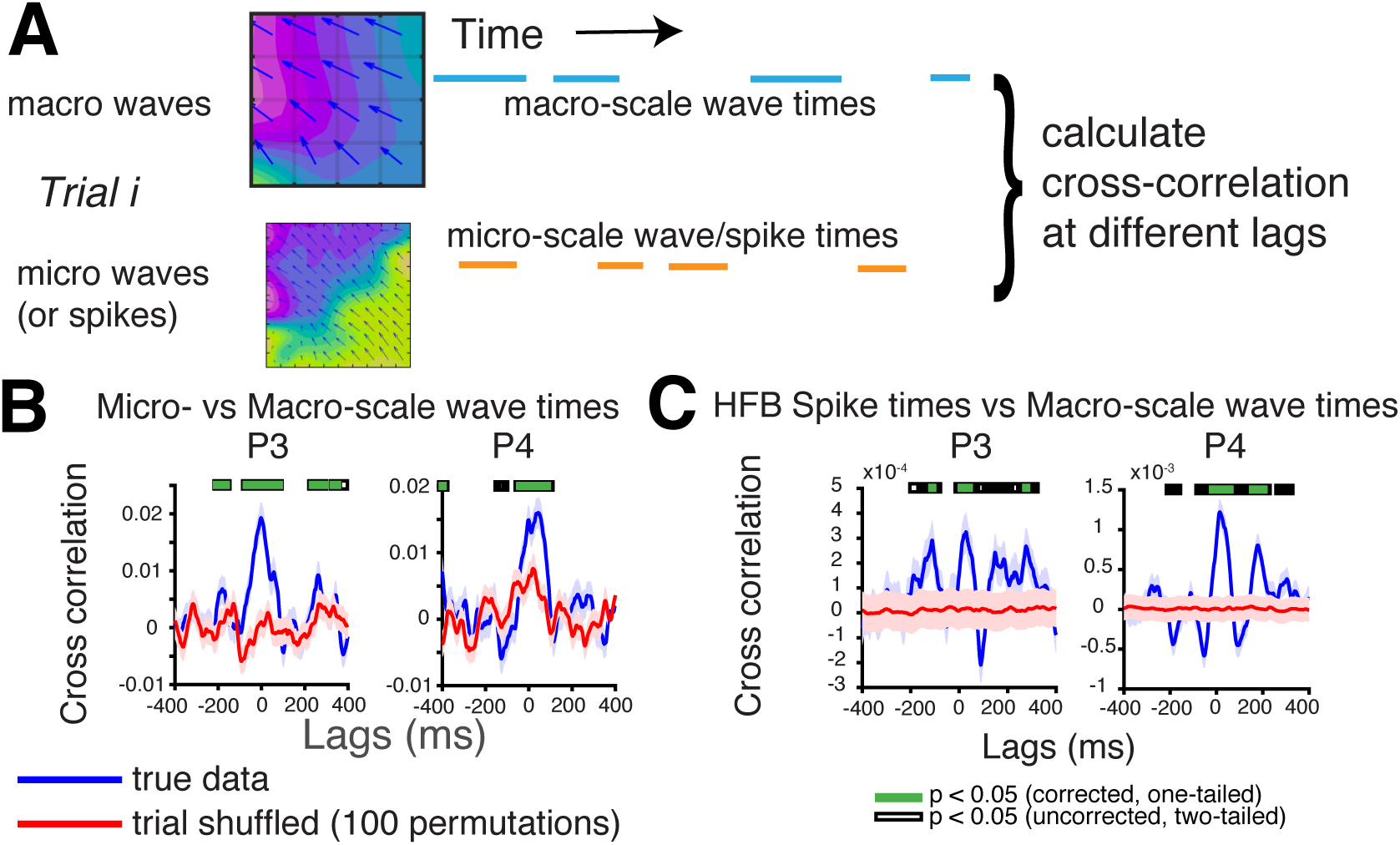
Relation between macro-scale traveling waves and micro-scale traveling waves and spiking activity. (*A*) A schematic of the analysis. We calculated the cross-correlation between the time series representing the time points during which a macro-scale (blue) or a micro-scale (orange) traveling wave was detected. We similarly computed the cross-correlation between the macro-scale wave time series and the time series of single unit spiking activity. (*B*) The cross-correlogram between micro-scale wave times and macro-scale wave times for two example participants. (*C*) The cross-correlogram between HFB spike times and macro-scale wave times for two example participants.

Given the temporal relation between macro- and micro-scale waves demonstrated by these data, we next investigated whether spiking activity is also temporally related to macro-scale traveling waves. We computed the cross-correlation between macro-scale waves and spikes as before, and compared the true correlograms to a distribution of shuffled correlograms (see Methods). We found a clear peak in the correlogram capturing the temporal relation between single unit spiking and macro-scale traveling waves when compared to the shuffled distribution in two participants (corrected *p <* 0.05; participants P2 and P6 in Supplementary Fig. S6C). One of these participants had a relatively high number of single units (P6). Data from the other participant with clean recordings and a high unit yield (P4) also indicated a relationship between macro-scale waves and spiking but this was not statistically significant. However, using HFB proxies for single unit firing, this relationship was clear in a majority of participants (Supplementary Fig. S6D and Fig. 5C), including both P4 and P6. We observed a greater peak in the average cross-correlogram between HFB spike times and macro-scale wave times at around time = 0 in four out of six participants in the true data compared to surrogate data (corrected *p <* 0.05), suggesting that low-frequency macro-scale waves are temporally related to spiking activity recorded along their path of propagation.

### Frequency-specific waves and memory processing

While we did not observe a relationship between 4 *−* 14 Hz waves and the memory task, it is possible that our broad filter that was designed with the explicit goal of maximizing temporal precision at the expense of frequency precision may have missed potential frequency-specific effects of traveling waves on behavior. Therefore, in order to further investigate the relationship between macro-scale waves and behavior, we reanalyzed the data using filters that were more precise in frequency. This reanalysis was performed by first filtering the iEEG and LFP data around each individual’s coherence peak (see Methods) and then recomputing optic flow and wave characteristics as before.

These frequency-specific waves also propagated in similar directions as the broad 4*−* 14 Hz waves described earlier and preferred to propagate in a posterior-anterior direction (Fig. 6A). Again, we observed that frequency-specific waves on the anterior temporal lobe (ATL) were more directional than those on the posterior (an example participant is shown in Fig. 6B; *t*(18) = 3.93, *p* = 0.001).

**Figure 6.**
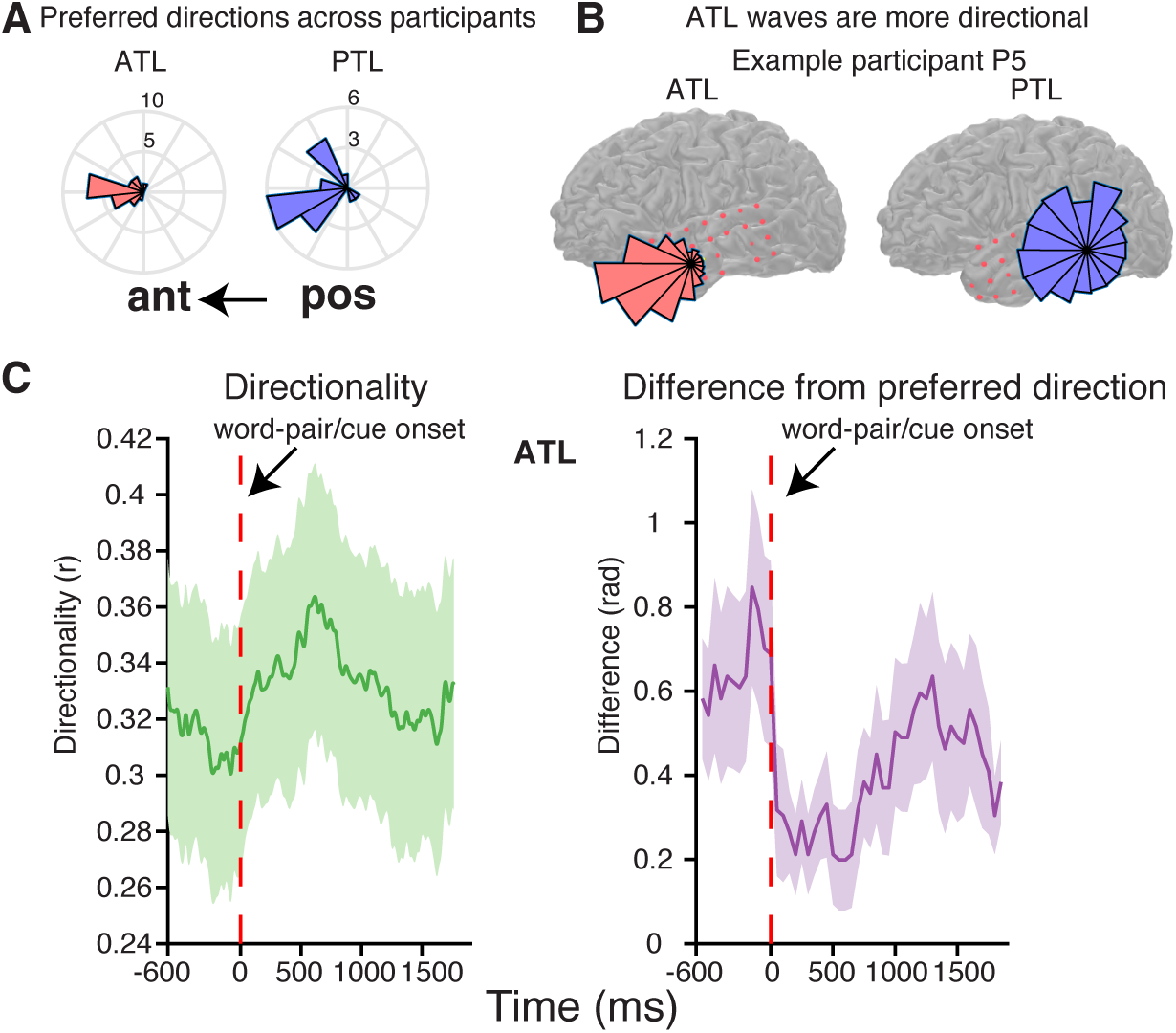
Frequency-specific macro-scale traveling waves on 4 *×* 4 subgrids and their relationship with the memory task. (*A*) The preferred directions of frequency-specific traveling waves in the anterior temporal lobe (ATL) and posterior temporal lobe (PTL) across all participants, showing a posterior-anterior preference. (*B*) Direction distributions for an example participant showing that frequency-specific macro-scale waves in the ATL are more directional than those observed on the PTL. (*C*) Directionality of ATL frequency-specific macro-scale waves averaged across all participants (left panel) and the difference in direction from the overall preferred direction in 200 ms sliding windows, averaged across all participants (right panel). The shaded regions show SEM.

Critically, on each of the 4 *×* 4 subgrids, we observed a decrease in wave directionality from fixation to word/cue onset, and then an increase in directionality during memory processing (ATL wave directionality is shown in the left panel of Fig. 6C). We computed the DC (directionality consistency) trend as described in Zhang et al. (2018) (see Methods) and observed a difference between DC trend during the 600 ms baseline period compared to an equivalent period during memory processing right after the onset of the word-pair or cue on the screen (statistically significant in the anterior and posterior subgrids for the frequency-specific waves, *p <* 0.032, n.s. in the middle subgrid (*p* = 0.18) and n.s. for 4 *−* 14 Hz waves in the ATL (*t*(19) = *−*0.57*, p* = 0.58)).

In order to understand whether the increased directionality during memory processing corresponded to waves propagating in their preferred directions or a different direction, we calculated the difference between the mode direction in 200 ms time bins and the overall preferred direction (see Methods). We performed this analysis with the ATL frequency-specific waves which were more directional than waves on other electrode subgrids and found that the increased directionality during memory processing corresponded to waves propagating closer to their preferred direction (right panel in Fig. 6C; see Methods). There was a significant difference between the deviation of waves from their preferred direction before word/cue onset and 500 ms after word/cue onset (*t*(18) = 3.30*, p* = 0.004; n.s. for the 4 *−* 14 Hz waves (*t*(19) = 0.52*, p* = 0.61), see Supplementary Fig. S7A,B and Supplemental Information to compare wave occurrence/direction changes with task variables between frequency-specific waves and broadband 4 *−* 14 Hz waves). This alignment with the preferred direction could not be explained by a difference in probability of wave occurrence (comparing the same time periods, *t*(18) = 1.3160*, p* = 0.21).

## Discussion

Our data demonstrate a direct link between traveling waves at the macro-scale, traveling waves at the micro-scale, and neuronal spiking activity. Although macro-scale and micro-scale waves appear to be temporally coordinated, our data suggest that they are not the same phenomenon. Micro-scale waves propagate along a different direction and with a slower speed than macro-scale waves. Critically, spiking activity is tightly linked to the propagation of micro-scale traveling waves. As micro-scale waves are temporally coordinated with macro-scale waves, it follows that macro-scale waves should also be related to spiking activity and indeed our data show that macro-scale waves are also closely linked to spiking activity. Both intra-regional and inter-regional correlations between spiking activity are important for cognition (Campo et al., 2015). Our results, made possible by the use of concurrent recordings from three different spatial scales in the human brain, suggest that traveling waves may play a role in cognition by coordinating spiking activity both across different brain regions and within local cortical regions.

The link between traveling waves at two different spatial scales and neuronal spiking suggests that traveling waves may provide a spatiotemporal framework for neuronal firing, and consequently information processing, in the human brain. For example, lower frequency traveling waves may underlie how spiking activity propagates across the cortex during individual tasks (Wu et al., 2008; Zhang et al., 2018). As high frequency activity is often taken as a surrogate for underlying local spiking activity (Manning et al., 2009), such coordinated activity is consistent with studies demonstrating the emergence of cross-frequency coupling during behavior (Axmacher et al., 2010; Canolty et al., 2006; Mormann et al., 2005). In addition, the presence of traveling waves that coordinate local spiking activity is also consistent with theoretical accounts regarding communication between brain regions which propose that coherence between low frequency oscillations in different brain regions underlies communication and coordination of spiking activity (Chapeton et al., 2019; Fries, 2005; Haegens et al., 2011). Traveling waves may be one mode of activity in the brain that facilitates such coherence.

If traveling waves therefore participate in coordinating neural processing across brain regions, it is not surprising then that we found traveling waves that were ubiquitous at both the macro- and micro-scales. The awake brain is always active, processing external stimuli or generating internal thought. Even during trials in which participants fail to successfully encode or retrieve a particular memory, they are likely engaged in some cognitive processing. The ubiquitous nature of traveling waves and their possible general role in coordinating information between brain regions may therefore explain why it has been difficult to identify such clear differences in traveling waves related to memory.

Our data suggest that traveling waves around the peak coherence frequency may play a role in memory formation and retrieval by becoming more directional and moving more consistently along their overall preferred directions (typically posterior-anterior across the temporal lobe) during memory processing. However, we observed no synchronization of the waves to the onset of word pairs or the memory probe in the task. During processing of higher-order conceptual associations, neural activity is more complex than that observed in anesthetized animals presented with visual grating stimuli. Interactions between different types of propagating wave patterns may lead to variable times at which coherent plane propagating patterns are detected, especially in the anterior part of the brain, away from the early visual regions. The specific role that traveling waves play in the service of successful memory processing still requires further exploration even though both our data and that of Zhang and colleagues (Zhang et al., 2018) suggest that directionality is likely to be the key feature of traveling waves related to cognitive task performance. On a related note, we included a technical note in the Methods section exploring the influence of choice of filters on estimated durations of waves detected using phase-based methods. It is possible that some existing reports of behavioral effects were partly driven by filtering choices and the time-frequency tradeoff that spuriously extended waves in time when spectrally highly precise filters were used. The raw and filtered traces in Fig. 1E show that our 4*−* 14 Hz filters do not temporally extend any of the oscillations in the data but more importantly, indicate that while there may be oscillations present in individual channels, neural activity over a large spatial extent does not typically align in a phase-lagged manner for much longer than a single cycle of the oscillation. While our study focused on such spatially extended propagating patterns of neural activity, other studies have focused on smaller collections of electrodes that exhibit similar patterns that may indeed extend for longer durations. However again, the technical discussion we presented in the Methods section suggests strongly that any traditional spectral analysis-based account of propagating waves must take into account the influence of the time-frequency resolution trade-off on the results. The transient waves we observed may also be partly explained by local coherent wave patterns interacting with each other. If such interactions are common across the brain, then it would be rare to see oscillations that line up across large extents of space in a time-delayed fashion, extending across several cycles of the oscillation. In fact, such interactions are thought to underlie distributed and parallel dynamic computation in the brain (Gong and Leeuwen, 2009). Therefore, our observation of transient wave-like patterns is consistent with the idea that there are local coherent propagating waves in the brain that interact with each other. In addition to the recently suggested possibility of interactions between local wave-patterns underlying dynamic computation, there are early suggestions that traveling waves cause sensory regions to mimic a “bar-code scanner” such that a fraction of the sensory field elicits an optimal response at each point in time (Ermentrout and Kleinfeld, 2001). Furthermore, Ermentrout and Kleinfeld (2001) also speculated that the presence of traveling waves indicates that neuronal activity in different spatial locations can be tagged with different phases of the oscillation, thereby providing a potential mechanism for categorizing and segmenting different elements present in the sensory input. One piece of evidence they cite for the plausibility of the idea is that the total variation in phase across the spatial extent covered by traveling waves is typically less than 2*π*. Given the variability in speed and directions of traveling waves, if the temporal extent of these waves significantly exceeds the duration of a cycle of the oscillation, then it is possible that different phases get tagged with the same sensory elements across different sweeps of the wave, making this scheme inefficient. However, our results indicate that low frequency traveling waves in the human cortex typically do not extend much longer than a cycle of the oscillation, making Ermentrout and Kleinfeld (2001)’s phase-tagging mechanism plausible.

Although our data demonstrate that traveling waves at the macro- and micro-scales are temporally coordinated, they also suggest that the waves at the two different spatial scales are not the same phenomenon. Waves captured using the micro-scale LFP recordings exhibit slower speeds and propagate in a different direction than the macro-scale waves captured through the iEEG contacts. This distinction in velocity between the two spatial scales is consistent with prior observations (Muller et al., 2018). For example, the slower propagation speeds we observed at the micro-scale are similar to prior reports of slower speeds in the human motor cortex at a similar spatial scale (e.g. 0.1 m/s in the 15 *−* 30 Hz range) (Takahashi et al., 2011). Similarly, the faster speeds at the macro-scale are consistent with previous studies of traveling waves using iEEG recordings (Zhang et al., 2018). The differences observed between the dynamics of waves at the two different spatial scales suggest that distinct axonal fiber tracts carry waves at the two scales (Muller et al., 2018). Specifically, the faster macro-scale speeds are more consistent with axonal conduction speeds of myelinated cortical white matter fibers whereas the slower micro-scale speeds are more consistent with axonal conduction speeds of unmyelinated long-range horizontal fibers within the superficial layers of the cortex (Muller et al., 2018). Therefore, while it may initially seem counterintuitive that waves that co-occur at the two different spatial scales propagate in different directions at different speeds, one possible explanation is that waves at the micro-scale are governed by the architecture of local neural circuits (Fig.3B), which may impose different constraints on the propagation velocity and direction of traveling waves than the constraints that govern larger scale traveling waves between brain regions. Yet despite these differences, we also found that traveling waves at the micro-scale were temporally coordinated with waves at the macro-scale. One possible mechanism by which this coordination could occur is via large-scale waves of depolarization that increase the probability of spiking (Wu et al., 2008), which in turn could trigger waves at the micro-scale (Nauhaus et al., 2009). These micro-scale waves may then have a further depolarizing effect recruiting more neurons to fire as they propagate, which would explain why the probability of spiking increases further after the onset of micro-scale waves in our data.

A similar coordination between traveling waves at two different spatial scales in humans occurs during epileptic seizures (Smith et al., 2016) where the propagating edge of the seizing territory (i.e., the ictal wavefront) is the source of traveling waves of ictal discharges that propagate out into the local cortical area at the micro-scale. These micro-scale traveling waves in turn depolarize the local cortical region and recruit more neuronal firing, consistent with our observation of micro-scale wave-locked spiking increases. In these hypersynchronous brain states however, the directions of ictal discharge propagation align across spatial scales prior to the recruitment of the MEA into the seizure, and the micro-scale directions of propagating discharges reverse after the passage of the ictal wavefront over the MEA zone. These dynamics and patterns of traveling waves during hypersynchronous pathological states suggest that wave activity in the context of seizures are fundamentally different from the wave propagation we observe during the more physiological non-seizing states we observe here.

The traveling waves we observed predominantly traveled from a posterior to anterior direction along the lateral temporal lobe. This direction of propagation at the macro-scale is also consistent with previous reports of traveling waves across the human cortex (Zhang et al., 2018). Importantly, we used an optic flow algorithm to automatically detect time periods of traveling wave activity. This approach detects and classifies traveling waves on a millisecond by millisecond basis, thereby allowing us to characterize the precise temporal locking of spatiotemporal activity across spatial scales and with underlying neuronal spiking. Our algorithm, however, requires the lagged-phase relationship between adjacent spatial sites to exist across the entire electrode grid. Hence, the algorithm only detects a traveling wave when most of the activity in the underlying cortical regions participates in the spatial phase gradient. It is possible, therefore, that additional traveling waves that involve only a subset of these larger cortical regions may be present but undetected by our algorithm. Similarly, it is also possible that even the identified traveling waves may have longer temporal durations but only involving a smaller region of cortex. Due to the constraints of our algorithm, these would also not be detected. Developing approaches that can identify these lagged-phase relationships with high spatial and temporal precision can help reconcile these issues. Finally, we did not measure eye movements in this study. It is possible that saccades trigger waves in the visual cortex (Zanos et al., 2015) which then propagate out over larger areas of the cortex and that any differences we observe between baseline and memory processing time periods may be driven by such saccades. Our stimuli were static word pairs but participants may have had to make large eye movements to read them. However, Zanos et al. (2015) reported that eye movements and visual stimuli were both needed to elicit traveling waves in the macaque V4, but that the directions of the waves were not affected by eye movements. The main characteristics of traveling waves that changed during memory processing were the directionality and directions which are unlikely to be affected by eye movements. Future work with eye-tracking can help tease apart the contributions of the stimuli themselves and the size of eye-movements to propagating waves observed in the human cortex.

Large-scale distributed cell assemblies are thought to be important for cognition. One mechanism by which these assemblies can be formed and maintained is via macro-scale waves that can spatiotemporally coordinate neuronal activity. Similarly, micro-scale traveling waves may recruit neuronal activity at a smaller spatial scale. While the direction of causality between spiking and waves is yet to be established, our data establishes the temporal relationships between traveling waves across different spatial scales and single unit spiking in the human brain that are necessary for such a role for traveling waves in neuronal communication. Therefore, traveling waves at multiple scales may underlie neuronal coordination and communication that is at the core of cognition and behavior.

## Materials and Methods

### Key resources table

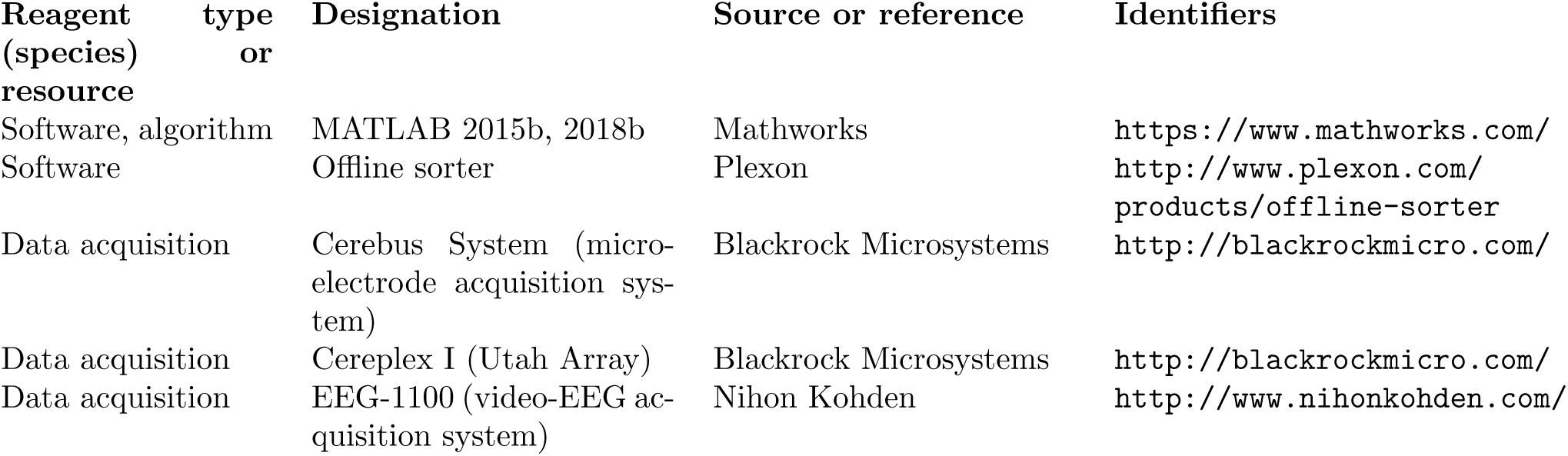

### Lead Contact and Materials Availability

Further information and requests for resources may be directed to, and will be fulfilled by, the Lead Contact Kareem Zaghloul.

### Experimental Model and Subject Details

20 participants (9 males, age 35.7 ± 2.3 years) with drug resistant epilepsy underwent a surgical procedure in which platinum recording contacts were implanted on the cortical surface. The clinical team determined the placement of the contacts in each participant so as to best localize epileptogenic regions. Data were collected at the Clinical Center at the National Institutes of Health (NIH; Bethesda, MD). The research protocol was approved by the NIH Institutional Review Board, and informed consent was obtained from the participants and their guardians. All testing was performed in the participants’ rooms during their stay in the Epilepsy Monitoring Unit. All analyses were performed using custom code written in Matlab (Natick, MA). We report standard errors of means unless otherwise specified.

### Method Details

#### Paired associates task

Each participant performed a paired associates task (Fig. 1B). Participants studied lists of pairs of words and later were cued by one word of the pair and had to remember the associated word they had learned earlier. Each participant performed one of two versions of the task that had slight differences in the experimental details as described below. As the tasks did not differ in the fundamental objectives and performance was indistinguishable between groups, we combined the data from both sets of tasks for subsequent analyses.

A single experimental session for each participant consisted of 15 or 25 lists, where each list contained either four or six pairs of common nouns shown on the center of a laptop screen, depending on whether the participant completed the first (*N* = 8 participants) or second version (*N* = 12 participants) of the task respectively. Words were chosen at random without replacement from a pool of high-frequency nouns and were presented sequentially and appeared in capital letters at the center of the screen. During the study period (encoding), each word pair was preceded by an orientation stimulus (either a ‘+’ or a row of capital X’s) that appeared on the screen for 250 *–* 300 ms followed by a blank interstimulus interval (ISI) between 500 *−* 750 ms (with a jitter of 75 ms). Word pairs were then presented stacked in the center of the screen for 2500 ms followed by a blank ISI of 1500 ms with a jitter of 75 ms in the first version of the task, or for 4000 ms followed by a blank ISI of 1000 ms in the second version. Following the presentation of the list of words pairs in the second version of the task, participants completed an arithmetic distractor task of the form A + B + C =for 20 seconds.

In both task versions, during the test period (retrieval), one word was randomly chosen from each of the presented pairs and presented in random order, and the participant was asked to recall the other word from the pair by vocalizing a response. Study word pairs were separated from their corresponding recall cue by a minimum lag of two study or test items. Each cue word was preceded by an orientation stimulus (a row of question marks) that appeared on the screen for 250 *−* 300 ms followed by a blank ISI of 500 *−* 750 ms. Cue words were then presented on the screen for 3000 ms followed by a blank period of 4500 ms in the first version of the task, or for 4000 ms followed by a blank period of 1000 ms in the second version. Participants could vocalize their response any time during the recall period after cue presentation. We manually designated each recorded response as correct, intrusion (when an incorrect word was vocalized), or pass. A response was designated as pass when no vocalization was made or when the participant vocalized the word ‘pass’. We defined all intrusion and pass trials as incorrect trials. A single experimental session therefore contained 60, 100, or 150 total word pairs, or study trials, depending on the task version and the same number of test trials, each cued by one of the two words in the pair.

#### Intracranial EEG (iEEG) Processing

We recorded intracranial EEG (iEEG) data from subdural electrodes (PMT Corporation, Chanhassen, MN) sampled at 1000 Hz using a Nihon Kohden (Irvine CA) EEG data acquisition system. Subdural macroelectrode contacts were arranged in both grid and strip configurations with an intercontact (center-center) spacing of 10 mm. The large scale traveling wave analyses in this study required an extended grid of electrodes. Therefore, we selected participants from a larger pool, with the requirement that each participant in this study has an 8 *×* 4 contact macro-electrode grid (Fig. 1A). Additionally, we only analyzed data from participants with fewer than 5 missing electrodes from the grid in order to get stable characterizations of traveling wave activity. Since the majority of these participants had temporal lobe grids, we restricted our analyses to the temporal lobe (*N* = 20 across both versions of the task). We filtered all iEEG traces using a notch filter (a fourth order Butterworth filter with a narrow 59 *−* 61 Hz stop band) to remove 60 Hz line noise.

We note here that since all behavioral and electrophysiological data come from participants with epilepsy, the results must be interpreted cautiously. In particular, it is possible that epileptic activity could contaminate our signals and may even manifest as the specific phenomenon we undertake to study here, i.e., traveling waves of activity. However, we took several steps to mitigate the effects of these confounds. First, we eliminated electrodes identified by the clinical teams during the monitoring period as being part of the ictal onset zone. Next, we removed trials that showed transient noise or epileptic activity using an automated algorithm tuned to match the assessment of the clinical team. The automated iterative procedure eliminates trials and channels that exhibited excessive variance or kurtosis (Wittig et al., 2018). We implemented this procedure by first extracting each trial’s data 600 ms before the word pair (for encoding) or cue (for retrieval) onset extending up to 4000 ms after onset, comprising a single trial. For each channel and trial, we computed the variance of the voltage trace during the 4.6 s surrounding the encoding or retrieval period. We used the resulting two-dimensional (trial *×* channel) matrix of variances to compute the maximum variance for each trial and channel. We calculated a threshold based on the quartiles of the resultant distribution of trial and channel variances as follows: 3*^rd^quartile* + *w_thresh_ × IQR*, where *IQR* is the interquartile range and *w_thresh_* is a user-specified parameter. If any trials exceeded the threshold, the maximum variance trials were iteratively removed and the quartiles were recomputed at each step. If any of the removed trials caused a change in the number of channels that exceeded the threshold, that trial was flagged for exclusion. Otherwise, if any channels exceeded threshold after all noisy trials were removed, those noisy trials were put back into the matrix and instead noisy channels were iteratively removed. This procedure of identifying noisy trials, and then channels, was iterated until all trials and channels were within the threshold limits. We used a *w_thresh_* of 0.5 (approximately 2 s.d. from the mean) for excluding trials as this was found to detect timepoints that were judged as noise by the epileptologist on our clinical team. However, we used a more liberal criterion of *w_thresh_* = 2.3 for excluding channels because we wanted to keep as many channels as possible for the spatial analysis while being more strict about keeping clean trials for analysis. Across the twenty participants, 2.4 ± 0.5 electrodes (range = [0, 7]) out of a total possible 32 electrodes arranged in an 8 *×* 4 grid) were either detected by the clinical team as being in the ictal onset zone or flagged by this iterative algorithm as being noisy and were removed. We also removed 14.2 ± 1.5% of trials using this cleaning procedure.

After removing noisy channels and trials as described above, we performed a global common average referencing by subtracting a common average computed across all channels separately for each trial and timepoint. One practical reason for this choice of reference, as opposed to a bipolar referencing scheme, was to retain phase information at each individual channel for the spatial wave analysis. We filtered the referenced data using a finite impulse response (FIR) filter with the first passband frequency at 4 Hz and second passband frequency at 14 Hz. The first stopband frequency was 1 Hz and the second stopband frequency was 18 Hz and stopband attenuation was set at 0.01. This equiripple 4 *−* 14 Hz bandpass FIR filter was designed using the filter design assistant (*designfilt*) in Matlab’s Signal Processing Toolbox. We performed the Hilbert transform on each electrode’s filtered data to extract instantaneous phase values at each time point. This procedure provided a spatial map of instantaneous phase values at each time point.

#### A technical note about choice of filters and its influence on phase-based wave analysis

Different studies employing phase-based wave analysis techniques have used different filtering methods centered around different frequencies. For instance, Zhang et al. (Zhang et al., 2018) used a 3*−*Hz band Butterworth filter (order not specified) around a specific frequency of interest, Muller et al. Muller et al. (2014) used an 8th-order Butterworth filter in a 5 *−* 20 Hz range, and Townsend et al. (Townsend et al., 2015) used an 8th-order Butterworth 1 *−* 4 Hz Butterworth filter. In one iteration of our analysis, we used a second order Butterworth filter to detect traveling waves in a (*f_c_ −* 1) - (*f_c_* +1) Hz band centered at the peak coherence frequency *f_c_* in each participant. Based on this approach, we detected waves at the macro-scale that frequently had durations of several hundred milliseconds, significantly longer than the durations of the waves we detected using FIR filters around the same frequencies but using a broader frequency range. This difference in durations has to do with the well known time-frequency resolution tradeoff. Increasing precision in the frequency domain (despite our use of only a second order filter with a slower roll off in the stopband) results in greater spread, and therefore lower resolution, in the time domain. Hence, it is possible that the longer durations of waves reported in previous studies may be artificially inflated by the filtering approach that was applied.

In order to further investigate the effect of the time-frequency tradeoff on phase-based wave analysis, we filtered the micro-scale LFP data using two directly comparable Morlet wavelets centered at the peak coherence frequency of 7 Hz for one participant’s data with different numbers of cycles (wave number = 3 versus 5). We ran the wave detection algorithm separately using the two different filters on the microelectrode data from this participant. The width parameter (typically defined as “the number of cycles”) of a Morlet wavelet controls the time-frequency precision trade-off such that the lower the wave number, the greater the precision in time and lower the precision in frequency. The full-width at half-maximum (FWHM) may be a more transparent way to quantify the width parameter (Cohen, 2019). The FWHM in the time domain of a 7 Hz Morlet wavelet with wave number = 3 is 160 ms whereas the FWHM for wave number = 5 is 268 ms. In the frequency domain, the FWHM is 4 Hz for wave number = 3 and 2.5 Hz for wave number = 5. We found that the average duration of micro-scale waves detected using wave number = 3 was 39 ms, whereas the average duration was 55 ms using wave number = 5 (two-sample *t*(15051) = *−*26.3, *p* = 5.1 *×* 10*^−^*^149^). This difference is due to the greater temporal spread that results from using the higher wave number.

To examine the effects of filter choice on the observed locking between waves and spiking activity, we computed the micro-scale wave-locked spike rasters using a spectrally precise (relative to the filter used in the main analysis) second order Butterworth filter in a 2*−*Hz band around an individual participant’s coherence peak frequency. The temporally precise relationships observed in the main analysis using a relatively spectrally imprecise 4 *−* 14 Hz filter were lost when using the 2*−*Hz band Butterworth filter (6*−* 8 Hz). The observed locking, however, could be recovered using a more spectrally imprecise (FWHM = 4 Hz) Morlet wavelet at 7 Hz (see Supplementary Fig. S8A).

While all the results above follow known time-frequency resolution tradeoffs, we decided to include this note to encourage future phase-based wave studies to pay careful attention to choice of filter depending on the goal of the investigation. Since our goal was to identify precise temporal relationships between phenomena at different spatial scales, we chose a broad low frequency band of 4 *−* 14 Hz that encompassed all individuals’ spectral coherence peaks for all the temporal coordination analyses presented in the paper. Furthermore, the results presented here also suggest that the choice of filters in phase-based propagating wave analyses may influence wave parameters such as duration which may potentially even have downstream effects on behavioral analyses.

#### Spectral coherence

We computed the magnitude squared spectral coherence between every electrode pair using the time series data from each electrode during each block. The coherence between two time series, *x*(*t*) and *y*(*t*), is a function of frequency:

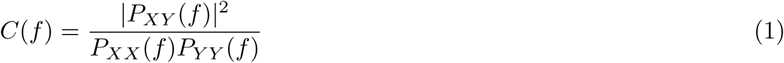

where *P_XX_* and *P_Y Y_* are the power spectral densities and *P_XY_* is the cross-spectral density. We estimated these spectral densities using Welch’s overlapped averaged periodogram method (Matlab function *mscohere*) (Cohen, 2014; Mitra and Bokil, 2009). Briefly, this involves dividing the time series into overlapping two-second epochs (50% overlap), tapering each epoch with a Hamming window, and then calculating the magnitude-squared spectral coherence for each epoch at each frequency from 2 to 100 Hz in steps of 2 Hz. The result is a single coherence spectrum for each electrode pair within each participant. We averaged the coherence spectrum over electrode pairs and we determined the frequency at which the spectral coherence peaked in each individual participant. We did this separately at the macro-scale and at the micro-scale.

#### Local field potentials (LFPs) and single unit spiking activity recorded by microelectrode arrays

We analyzed a previously reported dataset (Jang et al., 2017) to understand the relationship between the spatiotemporal patterns of traveling wave activity on the larger scale and the dynamic patterns of micro-scale local field potential and single unit activity on the smaller scale. Six participants (4 male; age 34.8 ± 11.5 years) were implanted with one or two 96-channel microelectrode arrays (MEAs, 4 *×* 4 mm, Cereplex I; Blackrock Microsystems, Salt Lake City, UT, USA) in the middle temporal gyrus (MTG) (see Fig. 1A for an example of placement of the MEA relative to the temporal grid electrodes). All six participants performed the second version of the paired associates task. MEAs were implanted only in participants with presurgical evaluation indicating seizure localization in the temporal lobe. Hence, the implant site in the MTG was chosen to fall within the expected resection area. Each MEA was placed in an area of cortex that appeared normal both on the pre-operative MRI and on visual inspection. Across participants, MEAs were implanted 14.6 ± 3.7 mm away from the closest macroelectrode with any ictal or interictal activity identified by the clinical team. MEAs arranged in a 10 *×* 10 grid structure had an inter-electrode spacing of 400 *µ*m and extended 1.5 mm into the cortical surface for five participants and 1 mm for one participant. Four out of the six participants received a surgical resection which includes the tissue where the MEAs were implanted. One participant had evidence of focal cortical seizure activity and received a localized resection posterior to the MEA site. One participant did not have a sufficient number of seizures during the monitoring period to justify a subsequent resection. Neither participant experienced a change in seizure type or frequency following the procedure, or experienced any noted change in cognitive function. The NIH Institutional Review Board (IRB) approved the research protocol, and we obtained informed consent from the participants explicitly for the placement of the MEAs and for all research components of this study.

We digitally recorded microelectrode signals at 30 kHz using a Cerebus acquisition system (Blackrock Microsystems, Inc.), with 16-bit precision and a range of ±8 mV. To obtain micro-local field potential (LFP) signals, we re-referenced each microelectrode’s raw voltage data by subtracting the average signal across all microelectrode channels, low-pass filtering the signal at 500 Hz, and downsampling at 1000 Hz. We used the same notch filter and automated iterative procedure described above for cleaning the iEEG data to remove line noise, and noisy trials and channels from the micro-LFP signals.

To extract neuronal spiking activity, we re-referenced each microelectrode’s signal offline by subtracting the mean signal of all the electrodes in the MEA, and then used a second order Butterworth filter to band pass the signal between 0.3 to 3 kHz. Using an offline spike-sorting software package (Plexon Offline Sorter, Inc., Dallas, TX, USA), we identified spike waveforms by manually setting a negative or positive voltage threshold depending on the direction of putative action potentials. The voltage threshold was set to include noise signals used in calculating unit isolation quality. Waveforms (duration, 1.067 ms; 32 samples per waveform) that crossed the voltage threshold were stored for spike sorting. Spike clusters were manually identified by viewing the first two principal components and the difference in peak-to-trough voltage of the waveforms. We manually drew a boundary around clusters of waveforms that were differentiable from noise throughout the experimental session. In this manner, we initially identified a total of 1469 putative single units.

Chronic extracellular electrodes have the potential to record from the same unit for multiple days. Therefore, one cannot assume that units recorded in different experimental sessions represent independent samples. To address this, we implemented a metric of unit identification to determine whether any units recorded across two consecutive experimental sessions were the same (Fraser and Schwartz, 2012). The metric assumes that distinct units can be separated using four characteristics of spiking activity: 1) waveform shape, 2) autocorrelation of the spike times, 3) mean firing rate, and 4) pairwise cross-correlation of spike times between units in the same recording session. Each pair of units recorded across two consecutive sessions was therefore represented by a single point in four-dimensional space using the four metrics above. Each four-dimensional similarity score, computed using a pair of units from different experimental sessions, can represent the similarity of either two units on different electrodes, which cannot be identical, or two units recorded from the same electrode, which may or may not be identical. We used a quadratic classifier to determine the decision boundary between different-electrode and same-electrode clusters in this four-dimensional space. This method identified 62 units that were putatively recorded for multiple sessions. The low number of overlapping units is expected, as sessions were on average at least two days apart (76 ± 19 hours), and because over such long periods, units may appear or disappear from recordings due to micromotion, changes in brain state, or other factors (Fraser and Schwartz, 2012). We removed these 62 units from all subsequent analyses to ensure independent samples across sessions.

Due to variability in the signal quality across recordings and the subjective nature of spike sorting, we quantified the quality of each unit by calculating an isolation score and signal to noise ratio (SNR) (Joshua et al., 2007). The isolation score quantifies the distance between the spike and noise clusters in a 32-dimensional space, where each dimension corresponds to a sample in the spike waveform. The spike cluster consisted of all waveforms that were classified as belonging to that unit, and the noise cluster consisted of all waveforms that crossed the threshold that were not classified as belonging to any unit. The isolation score is normalized to be between 0 and 1, and serves as a measure to compare the isolation quality of all units across all experimental sessions and participants. 1200 units (82%) had an isolation score of at least 0.9 which is the minimum threshold we used to include units in the analysis.

In addition to isolation quality, we computed the SNR for each unit using the following equation:

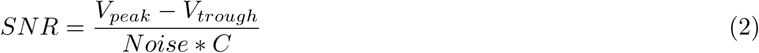

where *V_peak_* and *V_trough_* are the maximum and minimum voltage values of the mean waveform, and *C* is a scaling factor (set as 5) (Joshua et al., 2007). To obtain *Noise*, we subtracted the mean waveform from each individual waveform for each identified unit, concatenated these waveform residuals, and then computed the standard deviation of this long vector. Therefore, the noise term quantifies the within-unit variability in waveform shape. 1207 units (82%) had an SNR of at least 1.0. We additionally characterized the mean spike rate for each unit, and found that 914 units (62%) had a mean spike rate of at least 1 Hz. Overall, we retained a total of 633 units (43%) that had an isolation score of at least 0.9, a SNR of at least 1.0, and a mean spike rate of at least 1 Hz. The mean firing rate of all units was 3.24 ± 0.11, and the firing rate for the selected units was 4.24 ± 0.16 Hz.

To examine the temporal relation between micro-LFP traveling waves and spiking activity, we identified the starting time point of each micro-LFP traveling wave and collected spike trains that extended from *−*300 to +300 ms from this starting time point. We computed the sum of spikes at each millisecond over all micro-LFP waves across trials to get a summary picture of the micro-LFP wave-locked spike rasters in Fig. 4B and Supplementary Fig. S4A.

Since high quality single unit data could not be obtained for every participant, in order to facilitate comparisons across all participants, we constructed high frequency broadband (HFB) proxies for spiking. To do this, we first computed spectral power at 8 different log-spaced frequencies between 70 *−* 200 Hz using Morlet wavelets (number of cycles = 6) and then averaged the power across these frequencies. HFB proxies for spiking (henceforth called “HFB spikes”) were then constructed by identifying time points when HFB power *z*-scored within a microelectrode and trial exceeded *z* = 1.96. We confirmed that these HFB spikes reflected underlying single unit spiking by computing the cross-correlation between HFB spike events and single unit spike events in every participant (Supplementary Fig. S5A).

In order to examine whether spiking data may artificially confound the detection of traveling waves, we repeated our analysis after removing spikes from the LFP data (Supplementary Fig. S4B). For an example participant P4, we took the global average-referenced micro-local field potential (LFP) signal and identified the time points at which single unit spikes were detected on a given MEA channel. For each such time point, we interpolated the signal on that channel between *−*2 ms to +2 ms around the spike time, using a shape preserving piecewise cubic interpolation method to avoid sharp artifacts that may be introduced when using a linear interpolation (see Zanos et al. (2011) for a discussion of spike-removal methods). We used all sorted single unit data to select the time points for this interpolation, not filtering units based on SNR or isolation scores as we did for the main analysis.

#### Identification of traveling plane waves

Optical flow is defined to be the distribution of apparent velocities of movement of brightness patterns in an image (Horn and Schunck, 1981). For our purposes, we defined brightness at each electrode as the instantaneous phase value at that electrode. Optical flow therefore determines the velocities of movements of these phases across the electrode grid. We computed optical flow based on frame by frame spatial maps of instantaneous phase in a 4 *−* 14 Hz frequency band.

We used the opticFlow function from the MATLAB toolbox published by Townsend and Gong (Townsend and Gong, 2018) available at https://github.com/BrainDynamicsUSYD/NeuroPattToolbox to compute the velocity vector fields. The optic flow algorithm follows the Horn-Schunck method (Horn and Schunck, 1981). We provide a brief overview of the original approach here and refer readers to the original paper for further technical details (Horn and Schunck, 1981). Also see Afrashteh et al. (Afrashteh et al., 2017) for an overview of the different optical flow algorithms and comparisons of how well angle and velocity estimates match simulated ground truth. The Horn-Schunck algorithm works well for traveling plane waves (Afrashteh et al., 2017), which is the topic of our investigation here. A more recently described method performs slightly better when estimating speed (Bruhn et al., 2002). However, that method is a hierarchical algorithm that finds solutions from coarse to refined spatial scales that, in order to allow for spatial downsampling, requires a greater spatial coverage than we have access to in our data.

The first step in the Horn-Schunck method is to derive an equation to relate change in image brightness at a given point to the motion of the brightness pattern. The algorithm was developed based on two constraints. The first constraint, called brightness constancy, assumes that the brightness of a given pixel (*x, y*) is constant from one frame to next frame when it has moved to (*x* + *u, y* + *v*), leading to the equation:

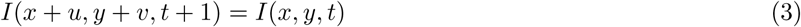

where *I*(*x, y, t*) is the brightness of pixel (*x, y*) at time t and *I*(*x* + *u, y* + *v, t* + 1) is the brightness of the same pixel when it has moved *u* in the *x* direction and *v* in the *y* direction in one time step (i.e., at time *t*+1). The Horn-Schunck method also assumes that the apparent velocity of the brightness pattern in an image varies smoothly across the image because if neighboring pixels all move in unrelated ways, there is little hope of recovering the velocities (Horn and Schunck, 1981). This leads to the second constraint, called spatial smoothness, which prevents discontinuities in the flow field. The spatial smoothness constraint is expressed as a minimization of the magnitude of the gradient of the optical flow velocity:

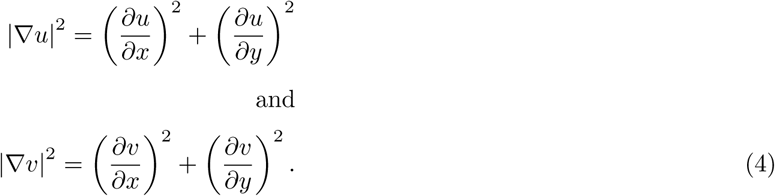

Both constraints are expressed together as a single minimization problem given by:

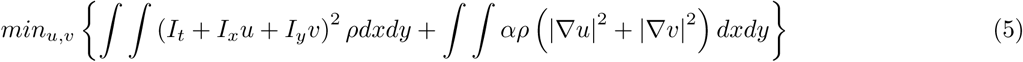

where *I_t_*, *I_x_*, and *I_y_* are derivatives of brightness *I*(*x, y, t*) with respect to time, direction *x*, and direction *y*. *α* is a smoothness regularization parameter that scales the global spatial smoothness term with respect to the brightness constancy term such that increasing *α* increases the influence of the spatial smoothness constraint on the solution for [*u, v*], producing a smoother velocity vector field. *ρ* is a penalty function. Horn and Schunck <u>(Horn and Schunck,</u> <u>1981)</u> used a quadratic penalty (*ρ*(*x*^2^) = *x*^2^). However, similar to Townsend and Gong (T<u>ownsend and Gong,</u> <u>2018),</u> we used the Charbonnier penalty, 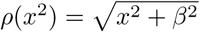, where *β* is a small positive constant. The Charbonnier penalty leads to more accurate velocity vector fields whereas the original quadratic penalty can lead to inaccuracies in the presence of hard edges (which are likely not an issue in neural data) or when adjacent regions move in different directions (which is certainly a possibility, for instance, when two concurrent traveling waves moving in different directions are present on adjacent regions of the temporal cortex). Equation 5 is solved by linearizing its corresponding Euler-Lagrange equations, to get a unique velocity vector field, [*u, v*] for each pixel (i.e., electrode) in the image (i.e., spatial map of instantaneous phase; see (T<u>ownsend and Gong,</u> <u>2018)</u> for more details). These velocity vectors therefore provide information about instantaneous speed as well as directions of motion of each pixel in the image.

The specific value of *α* and *β* for the optic flow model was determined by visually comparing the flow vectors obtained against movies of spatiotemporal changes in phase and choosing the values that provided the best visual match between the flow vectors and the apparent motion of activity in the phase movies (*α* = 1; *β* = 0.1). The resulting velocity vector fields were robust to small changes in these parameter values. These choices cannot be automated but Townsend and Gong (2018) suggest that small values of *β* result in a more robust non-linear penalty and result in more accurate velocity vector fields for regions with discontinuous motion which may be relevant for the macro-scale waves we characterized given that dynamical patterns may not smoothly vary across the entire temporal macroelectrode grid that spans several centimeters of the cortex. Townsend and Gong (2018) also suggest that reasonable values of *α* can range from *∼* 0.1 to *∼* 20 but that reduced spatial sampling (such as in the macro-scale data here) typically requires a smaller value of *α* to resolve dynamical patterns. Our observations and choice of parameters are consistent with Townsend and Gong (2018)’s recommendations.

A requirement for detecting traveling plane waves is that all vectors in the phase velocity field (PVF) point in the same direction. We used the average normalized phase velocity (Townsend et al., 2015):

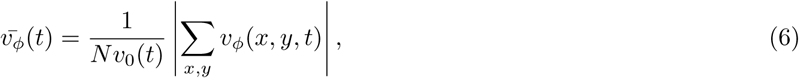

where *N* is the number of vectors in the PVF. For the macro-scale wave analysis, we split the 8 *×* 4 macroelectrode grid into three overlapping square 4*×*4 sub-grids (Figs. 1A,B) because of concerns that asymmetry in the geometry of the grid may cause the algorithm to have a directional bias simply due to the input 2D data having more information along one dimension than the other. This requirement of a minimum area of 3 *×* 3 cm^2^ captured by 4 *×* 4 macro-electrode sub-grids also eliminates possibilities such as neighboring locations happening to oscillate independently at a phase offset being erroneously called a traveling wave due to insufficient spatial sampling. Therefore, waves were detected on each sub-grid separately, and then eventually combined across the sub-grids to detect macro-scale waves that traveled across the entire 8 *×* 4 grid. Given that we identified an instantaneous phase at each electrode, where each electrode is analogous to each pixel in an image, this resulted in 16 velocity vectors for the macroelectrode 4 *×* 4 sub-grid data and 96 in the case of the microelectrode data arranged as 10 *×* 10 grids. *v*_0_ is the mean speed over all electrodes. Normalized phase velocity 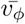 ranges from 0 to 1. 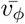 approaches 1 as all the vectors point in the same direction. We first imposed a threshold of 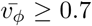 to detect all the timepoints where the vectors in the PVF align in the same direction. This by itself however does not capture waves traveling in the same direction across any extended duration because all the vectors may be aligned, but in different directions at different time points. Hence, this first step only flags long epochs during which sub-epochs contain velocity vectors that are aligned, but does not require that the velocity vectors identified in each sub-epoch be aligned in the same direction from one sub-epoch to the next.

In order to identify plane waves, we therefore imposed the additional constraint that plane waves must be comprised of uninterrupted timepoints during which the PVF points in the same direction. To identify such plane waves, we examined each identified long epoch that contained normalized phase velocities exceeding our threshold of 0.7. From each time point in each identified epoch, we calculated the distribution of directions in which the PVF pointed. To be considered a plane wave, we required that the PVF align with a fixed direction for a sufficient number of contiguous time points. We required the minimum time duration for a plane wave to be 15 ms. Given the possibility that the PVF may exhibit brief changes in direction due to noise, we considered a PVF as part of the same plane wave as long as the PVF returned to the same direction within 3 ms. Finally, to ensure that the PVF points in the same direction within each identified plane wave, we required that the mean directions across time points within that identified plane wave have a circular kurtosis of at least 0.7 and a median absolute deviation less than 15 degrees. These parameters were determined by trial and error, plotting movies of phase maps and ensuring that the detected wave epochs indeed look like traveling plane waves in the phase movies. All parameters were fixed before any of the other analyses reported in this study were performed.

Finally, after detecting traveling waves, we also applied a trial-level artifact rejection algorithm based on Staresina et al. (2015), also described in Vaz et al. (2019). Briefly, we applied a 250 Hz high pass filter in order to identify epileptogenic spikes and then calculated the gradient (first derivative) and amplitude at each time point. Any time point that exceeded a z-score of 6 in either the gradient or amplitude was marked as artifactual. If such an artefact was found on any of the 32 electrodes on the 8 *×* 4 grid, we removed any waves detected on any part of the grid during the artifactual time period in the trial. We also removed waves detected 30 ms before or after such artifactual epochs. We visually inspected the identified artifactual iEEG epochs and found that the automated procedure removed interictal epileptiform discharges (IEDs) and other artifacts as shown in the examples in Supplementary Fig. S8B. This procedure removed 11.8 ± 1.9% of the waves detected across subjects and subgrids. We observed an average of 2 electrodes on the 8 *×* 4 grid with artifacts per removed wave. Finally, we also examined the relationship between wave occurrence and epilepsy in the Supplementary Information (also see Supplementary Fig. S8C).

### Quantification and Statistical testing

#### Surrogate iEEG data

In order to compare the identified plane waves to any waves that would be identified by chance, we generated surrogate iEEG data with autocorrelations and spectral content that were similar to the original data, but with reduced cross-correlations between channels. Our goal was to generate surrogate data with reduced cross-correlations between electrode channels. We ran the wave detection algorithm on these surrogate data. If the surrogate data revealed a significantly reduced number of waves compared to the original data, then this would suggest that the traveling plane waves identified in the original data are indeed due to the specific phase-lagged relationship between spatial locations on the electrode array and are not an artifact of the analysis method itself.

To generate the surrogate iEEG data, we used the Prichard-Theiler method (Prichard and Theiler, 1994) which was also used as a control in a previous study of traveling waves (Townsend et al., 2015). The Prichard-Theiler method strives to preserve the cross-spectrum in multivariate data in addition to the power spectrum of each individual channel. We produced such phase-randomized surrogate data by multiplying the Fourier phases of each channel by a uniform random phase. Specifically, we added the same random phase, chosen from the uniform distribution 0 *−* 2*π*, to the phase values for each frequency component of the discrete Fourier transformed signal in each electrode. This approach preserves the cross-spectrum between channels since the cross-spectrum reflects only the relative phase differences between them. In addition, we shifted the phase at each frequency *f* by another random phase which was chosen from a normal distribution (mean = 0, SD = *π/*4) (Townsend et al., 2015). We applied this same random phase shift to the data from each electrode channel. This surrogate iEEG data thus produced preserves autocorrelations and power spectra but reduces the cross-correlation between channels by an average of 26.6 ± 0.6% (mean ± SEM) across participants (see Supplementary Fig. S1). The exact reduction in cross-correlation is not a parameter we control in this approach, but the procedure above resulted in a reduction of across-channel cross-correlations by approximately 25% which provided a reasonably conservative test.

#### Velocity calculation

The output of the optic flow computation provides both directions and magnitudes of flow. We verified the directions by visually examining movies of identified traveling waves superimposed with the flow fields but we observed that the algorithm tended to underestimate the magnitudes (or speeds). We found several discussions of the same issue faced by other users of the built-in MATLAB function *opticalFlowHS*. One possibility is that the algorithm due to the emphasis on minimization of gradients tends to underestimate spatial gradients. Indeed the original paper by Horn and Schunck (Horn and Schunck, 1981) mentions this possibility, noting that the velocity estimates “tend to be too small, rather than too large” (pp. 196).

For this reason, we implemented a method to compute velocities of the identified traveling waves, based on the method used in a recent publication on iEEG traveling waves (Zhang et al., 2018) (also see Rubino et al. (2006) for a similar method). We first computed the temporal frequency *f* of a traveling wave by first taking the average phase across electrodes at each time point and then calculating the temporal derivative of this average phase. This generates an estimate of instantaneous temporal frequency at each time point within the wave. We took the mean temporal frequency across time points to get *f* for each wave. We calculated the spatial frequency *ξ* by first taking a reference electrode and identifying the relative phases at all electrodes with respect to that reference phase. We then regressed the relative phases on Euclidean distance from the reference electrode to get an estimate of spatial frequency. We averaged this spatial frequency over all time points to get *ξ* for the wave. We did this directly on the time points known to exhibit phase progression across electrodes because the optic flow algorithm already *f* identified the time points of interest. We then estimated the wave speed as 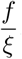. In order to get robust estimates at the macro-scale, we used long-range waves that propagated across the entire length of the grid of electrodes either in the anterior-posterior or the posterior-anterior direction (defined to be within 30 degrees on either side of the long axis of the grid) and those that lasted for at least 100 ms. We took the median speed within participants and took the mean across participants to summarize the speed of propagation.

#### Directionality calculation

Directionality (or directional consistency (DC)) of waves was defined to be the resultant vector length of all wave directions. We used the *circ r* function in the *CircStat* toolbox (Berens, 2020) to compute directionality. The trend in directional consistency (DC trend) was then computed by taking the Pearson correlation between directionality and time as in Zhang et al. (2018).

We computed the mode direction within each participant’s anterior temporal lobe (ATL) subgrid to characterize the “preferred” direction. The mode was calculated by taking 25 bins in a circular histogram and getting the mean value of the bin edges that capture the most number of waves. We confirmed visually that the obtained “preferred” direction was indeed the dominant direction seen in the overall distribution of wave directions for each individual and that this was robust across reasonable choices for the number of bins used in the mode computation. We then used time bins of 200 ms which were moved in steps of 50 ms to capture all waves across trials within a given participant and computed the mode direction within each time bin and finally computed the difference in angle between the mode direction in each time bin and the overall preferred direction. This procedure produced a single time course of angle differences from the preferred direction for each participant, and we then averaged these values across participants to obtain the average deviation from preferred wave direction which was plotted in the right panel of Fig. 6C. We compared the mean of the deviation from preferred direction in a baseline period of 200 ms right before word/cue onset with the mean in an equivalent period 500 ms after word/cue onset to test whether waves aligned better with their preferred direction during task processing. The observation of a fixation-related increase in frequency-specific waves (see Supplementary Fig. S7A) motivated the choice of this 200 ms baseline window which is 400 ms away from the time of fixation but still before word/cue onset.

#### Statistical testing

We performed statistical tests of circular distributions (e.g. directions or phases) on the identified waves using Hodges-Ajne tests when the null hypothesis was that the angles are uniformly distributed (e.g., where the alternative can be unimodal, bimodal, or anything other than uniform) and using Rayleigh’s tests when the alternative hypothesis was a preference for a specific angle (i.e., a unimodal distribution). We used the *CircStat* toolbox for MATLAB for directional statistics (Berens, 2020).

To examine the relation between traveling waves at the macro- and micro-spatial scales as well as between waves and spiking activity, we computed the cross-correlation between the vectors representing the time points at which traveling waves and spikes were detected. For example, in order to quantify the relation between macro- and micro-scale traveling waves, in each trial we constructed a binary vector that took on a value of 1 for each time point at which a traveling wave was identified at the macro-scale, and a separate binary vector identifying time points of traveling wave activity at the micro-scale. In order to examine the relation between macro-scale traveling waves and spiking, we followed a similar procedure using spike times and HFB spike times. For all pairs of phenomena, we then computed the cross-correlation between the two binary vectors to generate a cross-correlogram for each trial. A *t*-test was performed to determine the time points at which the true cross-correlogram across trials was significantly different from zero (black bars for uncorrected *p <* 0.05 in Supplementary Fig. S6). We then constructed surrogate data by permuting trial labels and recalculating the cross-correlograms in these permuted data and obtained *t*-statistics in each of 100 permutations. For each permutation, we recorded the maximum *t*-statistic across time lags and finally computed the proportion of permutations where the maximum *t*-statistic exceeded the true *t*-statistic at a given time point to get one-tailed p-values comparing the true and surrogate correlograms corrected for multiple comparisons across time lags (green bars for corrected *p <* 0.05 in Fig. S6).

### Data and Code Availability

Processed data of instantaneous phase maps and population spiking activity used in this study can be found at: https://neuroscience.nih.gov/ninds/zaghloul/downloads.html.

## Supplementary Information

### Macro-scale traveling waves and behavior

We were interested in whether and how traveling waves of neural activity may be related to behavior. In order to understand whether memory processing elicits waves, we first compared the likelihood of macro-scale waves (4 *−* 14 Hz) between two time periods while participants performed the paired associates task, a 0.6 s baseline period before the onset of the word-pair (at study) or cue word (at test) and a 0.6 s task period (to match the baseline period), starting 0.5 s after the onset of the word pair or cue in order to separate out the two periods. We found no significant differences between baseline and task periods in the likelihood of waves on any of the macro sub-grids (*t*(19) *> −*2.05*, p >* 0.054). Next, in order to understand whether macro-scale waves were related to successful memory processing, we considered traveling waves during a 4 s task period, starting 0.5 s after the onset of the word pair or cue. The likelihood of occurrence of traveling waves on the 4 *×* 4 subgrids did not differ between successful and unsuccessful encoding/retrieval time periods (*t*(19) *<* 1.22*, p >* 0.23).

We then examined long-range waves that propagated across all three sub-grids, and also observed no relationship between waves and successful associative memory processing. First, considering all long-range waves, we found no difference between the likelihood of waves in the successful memory encoding task periods compared to the unsuccessful memory encoding task periods (46.3 ± 1.8% versus 45.3 ± 2.0%; *t*(19) = 0.83*, p* = 0.42) nor between successful retrieval periods and unsuccessful retrieval periods (45.2±1.8% versus 44.7±1.8%; *t*(19) = 0.5965*, p* = 0.56). In order to obtain more robust waves for the behavioral analysis, we then considered long-range waves that lasted at least 71 ms, which is one cycle of the highest frequency in the frequency band that we used (14 Hz). Again, we found no differences between successful and unsuccessful memory task periods (encoding phase: 25.1 ± 1.8 versus 24.4 ± 1.6%; *t*(19) = 0.83*, p* = 0.42; retrieval phase: 23.9 ± 1.8% versus 24.5 ± 1.7%; *t*(19) = *−*0.77*, p* = 0.45). Finally, we split the long-range waves that lasted at least 71 ms into those that propagated from posterior to anterior and anterior to posterior. We did not find any differences between successful and unsuccessful memory conditions either during the encoding phase (19.1 ± 1.8% versus 18.8 ± 1.7%; *t*(19) = 0.24*, p* = 0.8) or the retrieval phase (19.0 ± 1.7% versus 18.6 ± 1.7%; *t*(19) = 0.55*, p* = 0.6) for the posterior-anterior waves. While the anterior-posterior waves were significantly less frequent than the posterior-anterior waves (see main Results), we observed a decrease in likelihood of these anterior-posterior waves during successful retrieval of words compared to unsuccessful retrieval (4.9 ± 0.1% versus 5.9 ± 0.1%; *t*(19) = *−*2.87*, p* = 0.01) but no differences between successful encoding and unsuccessful encoding periods (6.1 ± 0.1% versus 5.6 ± 0.1%; *t*(19) = 1.26*, p* = 0.22). Finally, we also examined long-range wave activity in the retrieval periods but locked to response times and observed no differences between successful and unsuccessful retrieval conditions at any point within a two second time window prior to vocalization (*p >* 0.05, two-tailed t-test, corrected for multiple comparisons by using the Benjamini-Hochberg procedure to control the false discovery rate (FDR)). In the incorrect condition, when participants did not vocalize a response, a reaction time was assigned to that trial by drawing randomly from the distribution of reaction times during correct retrieval.

While we did not find a relationship between 4 *−* 14 Hz waves and behavior, when looking at frequency-specific waves around the coherence peak, we observed clear changes in directionality after word/cue onset during the task (see main text and Results). Due to low memory success rates in many participants, we could not however discern any robust relationships between directionality and memory success because the circular mean of directions is sensitive to the number of samples. For instance, only 8 participants responded correctly to memory probes with probability greater than 0.3 and had at least 50 correct trials compared to 255 ± 42 incorrect trials across participants. However, these results indicate that traveling waves near the peak coherence frequency are modulated by the task. We also observed an increase in the probability of observing a long-range frequency-specific wave following the offset of the fixation cross (Supplementary Fig. S7A) but this was not seen earlier using the broadband low frequency waves. The probability of observing a long-range peak coherence-frequency wave in a 600 ms baseline period was significantly greater than in an equivalent period during memory processing (starting 500 ms after the words/cue appeared on the screen, in order to separate out the baseline and task periods; P(wave) = 0.13 versus 0.10; *t*(18) = 3.6313*, p* = 0.002) but this difference was driven by the increase observed right after the fixation cross at *t* = 0 in Supplementary Fig. S7A). There was no such difference between baseline and task periods observed using the broadband low frequency waves (*t*(19) = 0.2703*, p* = 0.79).

Our data suggest that frequency-specific traveling waves may play a role in cognition, however, we need to make methodological advances to address issues related to the time-frequency resolution tradeoffs we discussed in the Methods section (also see Discussion in the main paper).

### The effect of epilepsy on wave occurrence rates

In order to understand the role of epilepsy on the observed waves, we first calculated the correlation between the rate of wave occurrence on the anterior, middle, and posterior 4 *×* 4 subgrids and different measures of severity and duration of epilepsy across the 20 patients. Specifically, we calculated the correlation between wave occurrence rates and 1) age at surgery, 2) seizure duration (21.6 ± 2.4 years), and 3) seizure frequency (5 patients had daily seizures, 10 weekly, and 4 monthly with no information for 1 patient; Spearman’s rank correlation). For all subgrids, we found a significant correlation between seizure duration and wave occurrence rates (rho *≥* 0.48, *p ≤* 0.03; *p >* 0.56 for correlations with age at surgery and seizure frequency; Supplementary Fig. S8C).

We further compared “clean” waves on the anterior and posterior subgrids with the “unclean” ones detected within 30 ms of artifacts and IEDs anywhere on the 8 *×* 4 grid (see Methods). The directionality of waves (measured by the length of mean resultant vector of directions) did not differ significantly between clean and unclean waves (*t*(19) *<* 1.45*, p >* 0.17). However, the waves detected during artifacts on the grid were slightly but statistically significantly longer lasting than the clean waves (30 ± 1 ms vs 28 ± 1 ms on the anterior subgrid, *t*(19) = 4.96*, p* = 8.7 *×* 10*^−^*^5^; and 27 ± 1 ms vs 26 ± 1 ms on the posterior subgrid, *t*(19) = 4.63*, p* = 1.85 *×* 10*^−^*^4^).

These analyses suggest that the epileptic brain, by virtue of increased local and global connectivity, may show increased propagating wave activity that is related to the duration of epilepsy. The waves that co-occur with IEDs and artifacts however do not differ in directionality from the waves that are clean of IEDs, but they tend to be longer in duration. Finally, we provide more information about epilepsy onset, duration, seizure type, seizure frequency, and medication in Supplementary Table 2. Höller et al. (2018) provides an overview of pharmaco-EEG studies with anti-epileptic drugs (AEDs) based on 37 studies. Different AEDs had different effects on oscillatory activity. For example, Carbamazepine seems to increase theta and reduce alpha activity in most studies whereas Phenytoin and Lamotrigine seems to reduce theta and alpha oscillatory activity. In general, the review indicates a slowing of rhythmic activity in the EEG (increased activity up to 7 Hz and decreased activity in higher frequency ranges). Since the reported coordination between traveling waves at different spatial scales and spiking activity holds after extensive cleaning of IEDs and other artifacts, and given the observed change in directionality of frequency-specific traveling waves during memory processing, the main conclusions in the paper are unlikely to be driven by epilepsy.

## Acknowledgements

This work was supported by the Intramural Research Program of the National Institute for Neurological Disorders and Stroke. We are indebted to all patients who have selflessly volunteered their time to participate in this study. This work utilized the computational resources of the NIH HPC Biowulf cluster (http://hpc.nih.gov) and we thank Chris Zawora for help with running the analysis code on the supercomputer. We thank Alex Vaz and Zane Xie for helpful comments on the manuscript and Deepak Gopinath for proof-reading.

---

**Figure S1.**
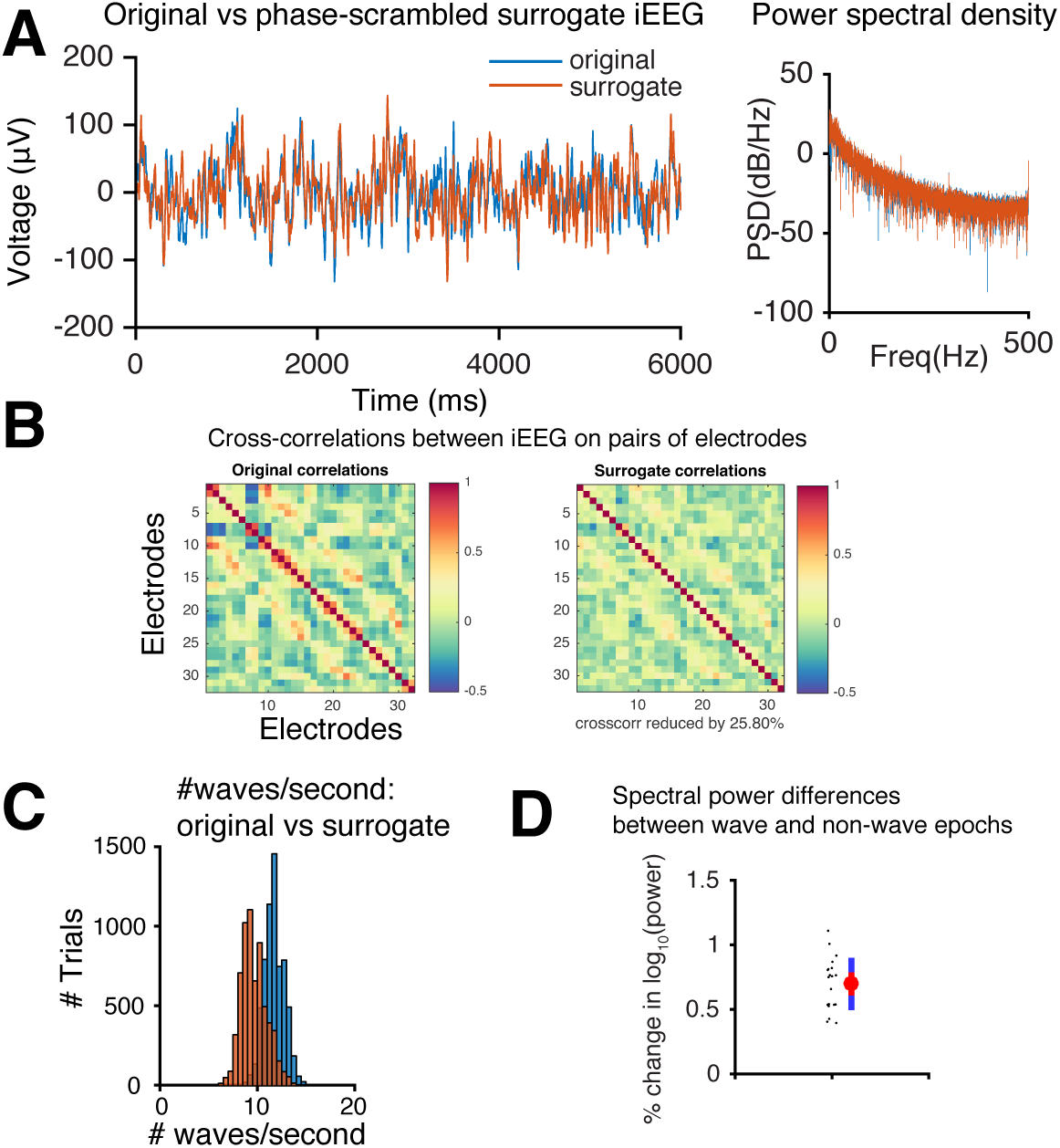
Surrogate data construction and comparison of waves detected in the real versus surrogate data. (*A*) An example voltage trace (blue) and the corresponding phase-scrambled surrogate trace (orange), illustrating the constructed surrogate that mimics the original trace very closely except for the phase information. The power spectral density for the original and surrogate data is shown on the right. (*B*) The spatial cross-correlations between voltage traces on pairs of electrodes (total 32 electrodes arranged as an 8 *×* 4 grid). The cross-correlations are decreased by around 27% in this example surrogate data constructed by the Prichard-Theiler method (see Methods). (*C*) The distribution of the number of waves per second detected in the original data compared with the number detected in the surrogate data. A clear and significant decrease is observed in the number of waves detected in the distorted surrogates (11.74 ± 0.10 versus 9.91 ± 0.23; *t*(19) = 8.93, *p* = 3.15 *×* 10*^−^*^8^). Greater distortions in the phase information that would reduce the spatial cross-correlations between electrodes would result in an even greater separation. (*D*) Percent change in log power from non-wave to wave epochs (*t*(19) = 21.92*, p* = 6 *×* 10*^−^*^15^).

**Figure S2.**
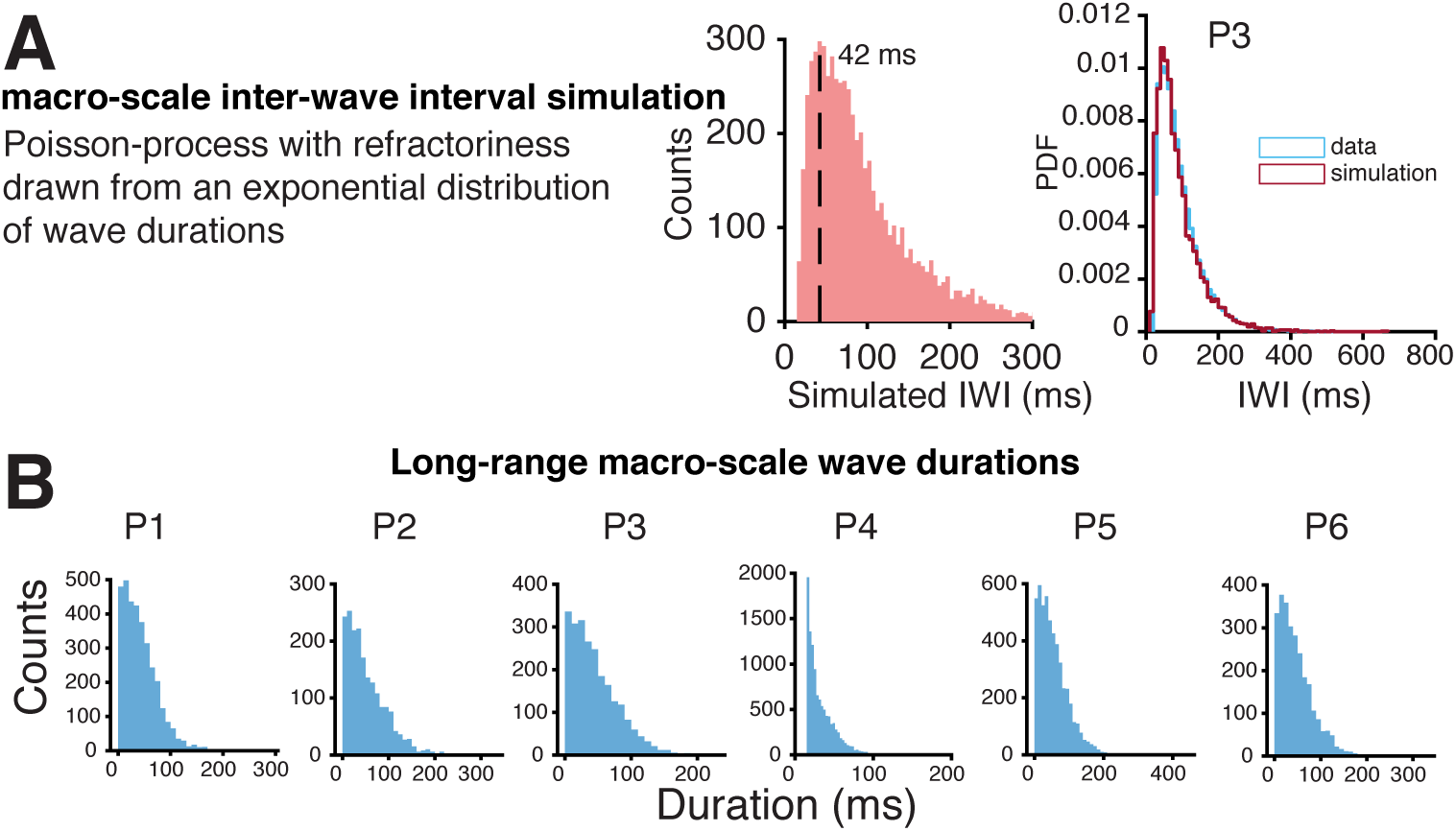
Simulated inter-wave interval distribution and distribution of durations of traveling waves at the macro-scale. (*A*) The distribution of macro-scale inter-wave intervals (IWIs) simulated by assuming a Poisson process with a refractory period that is drawn from an exponential distribution of wave durations and with a rate that is similar to the rate of occurrence of macro-scale (4 *×* 4 macro-subgrid) waves in our data of *∼* 11 per second. The shape of this distribution resembles that of the IWI distributions presented in Fig. 2D. An example participant P3’s IWI distribution and the simulated distribution overlaid on top is shown on the right. (*B*) The distributions of durations of long-range traveling waves in six example participants.

**Figure S3.**
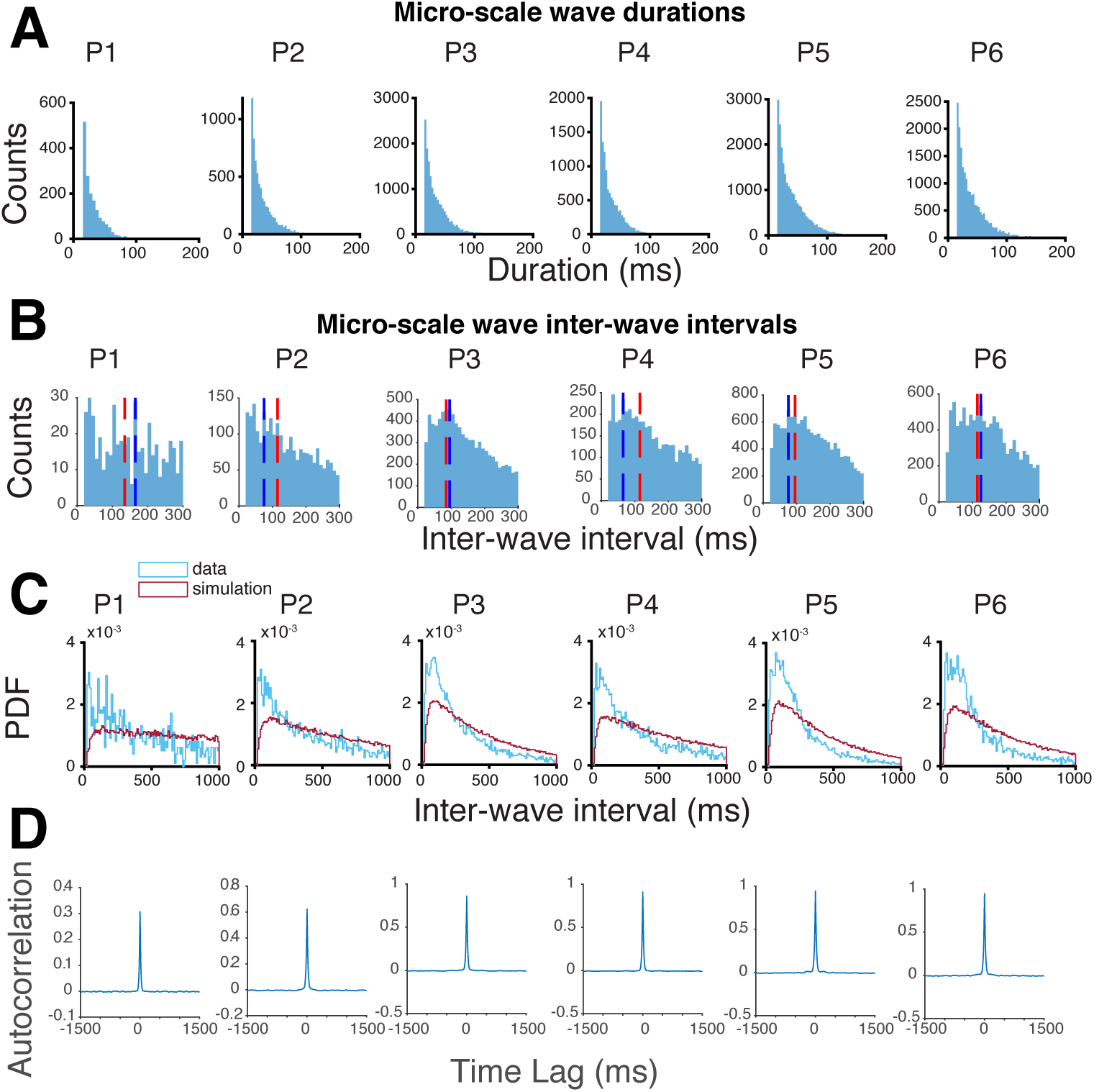
Simulated inter-wave interval distribution and distribution of durations of traveling waves at the micro-scale. (*A*) The distributions of durations of micro-scale traveling waves in all six participants with MEA recordings. (*B*) The distributions of inter-wave intervals (IWIs) of micro-scale traveling waves in all six participants with MEA recordings. Blue dotted lines indicate the empirical peaks away from time = 0 and red dotted lines mark the peaks in the simulated IWI distributions in *C*. (*C*) Micro-scale IWI distributions simulated using the same procedure as in Supplementary Fig. S2A but by choosing the rate parameter to match each individual’s measured rate of occurrence of micro-scale waves. Note that the X axis has been expanded to include time intervals up to 1000 ms. (*D*) The autocorrelations in micro-scale wave times.

**Figure S4.**
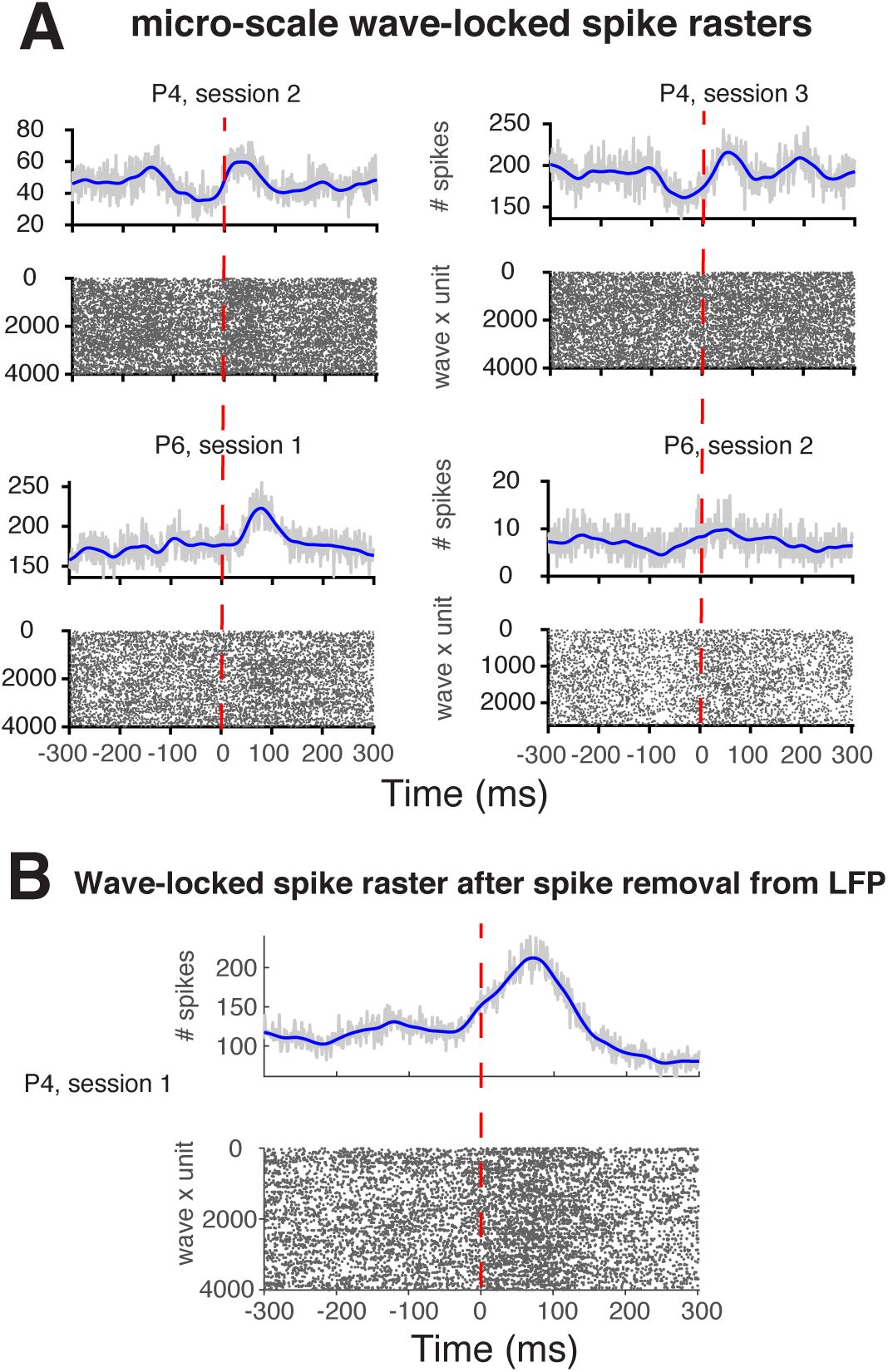
Examples of spike rasters locked to the onset of micro-scale traveling waves. (*A*) The onsets of micro-scale traveling waves are followed by increases in spiking activity. (*B*) Micro-scale wave-locked spike rasters for participant P4 after spike-removal from the LFP

**Figure S5.**
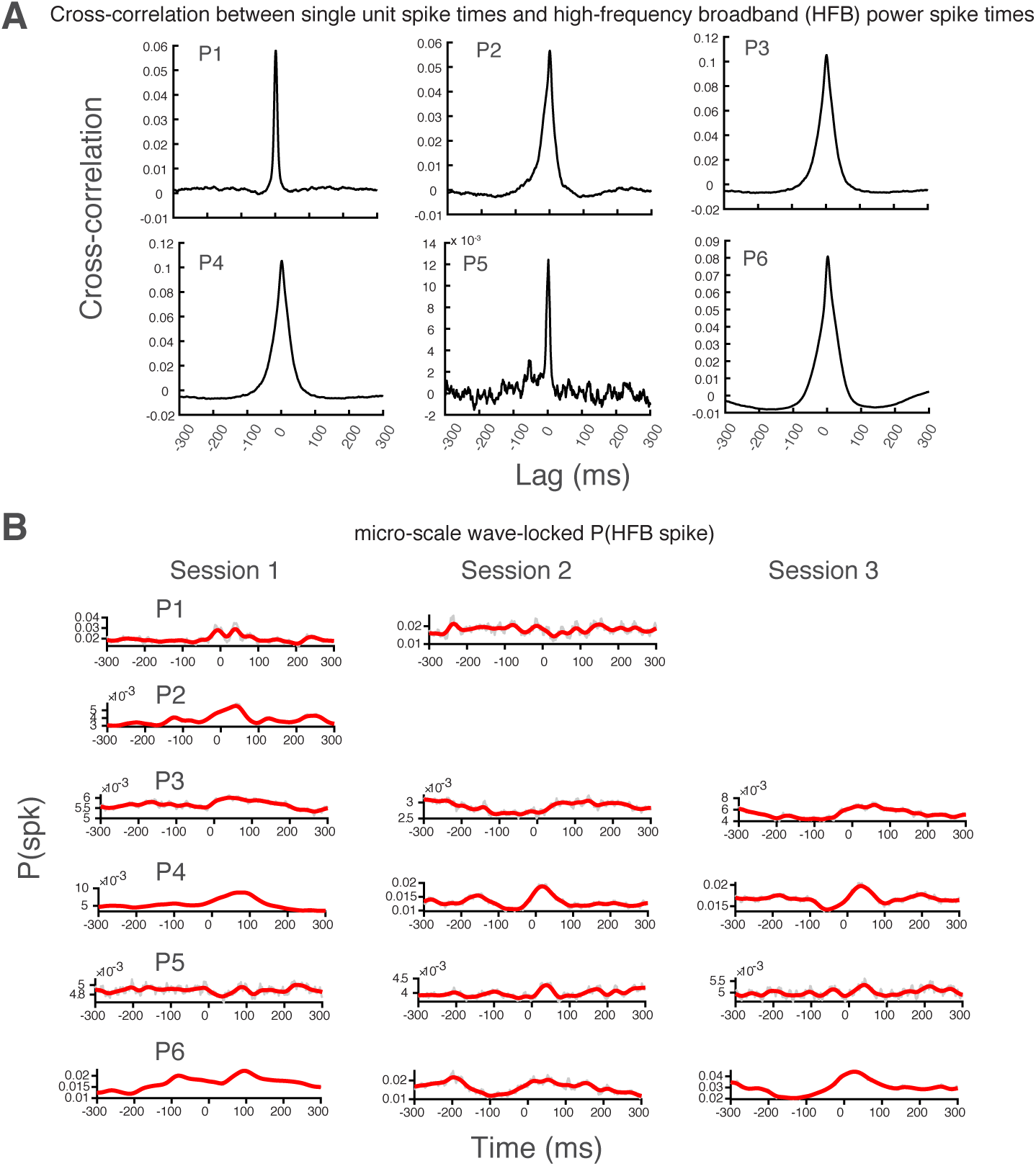
High-frequency broadband (HFB) proxy for spiking is locked to the onset of micro-scale traveling waves in all participants. (*A*) Cross-correlations between binarized continuous time high frequency broadband power and multiunit spiking activity in each participant (see Results). Single unit spike trains and HFB spike trains are cross-correlated at 300 different lags in both forward and backward directions within trials, and then averaged over trials and sessions. The correlograms peaked at time = 0 suggest that high frequency power can be used as a proxy for spiking activity. (*B*) We detected time points at which HFB power exceeded a threshold within each trial and micro-electrode and called these HFB spikes. Probability of detecting an HFB spike in each participant and in each session was computed as the proportion of HFB spikes at each millisecond across all waves and trials locked to the onset of micro-scale traveling waves (time = 0), and is plotted in gray. A smoothed version (using a Gaussian window of 50 ms) of the probability is overlaid in red.

**Figure S6.**
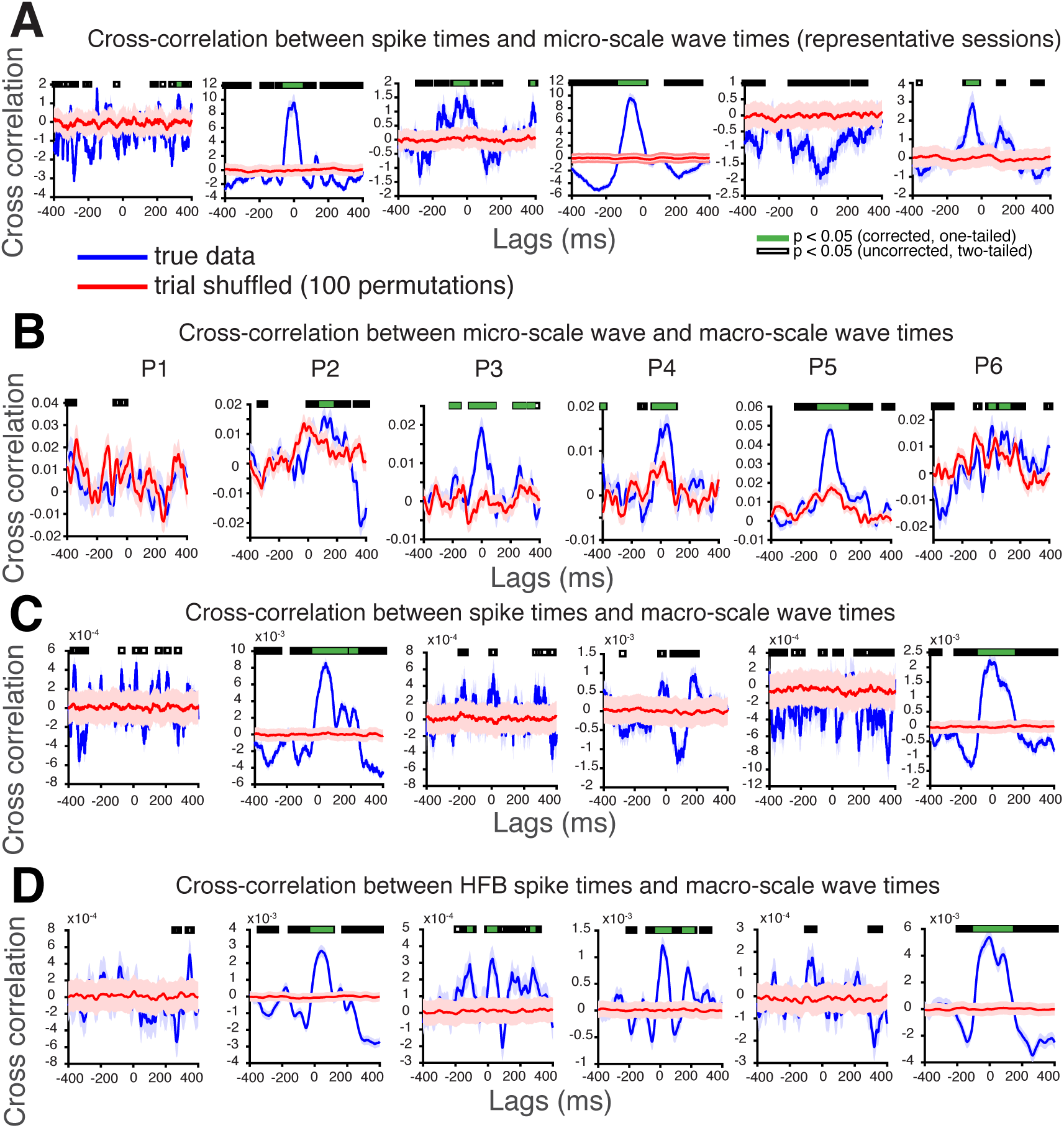
Cross-correlation between the time series of spiking activity, micro-scale waves, and macro-scale waves detected on the nearest macroelectrode subgrid to the microelectrode array. (*A*) Cross-correlation between the time series of single unit spiking and micro-scale wave times. A trial-shuffled distribution of cross correlations is shown in red (see Methods). (*B*) Cross-correlation between the time series of detected micro-scale and macro-scale traveling waves in each participant. (*C*) Cross-correlation between the time series of detected single unit spiking events and the time series of macro-scale waves. (*D*) Cross-correlation between the time series of HFB spiking events and macro-scale wave times.

**Figure S7.**
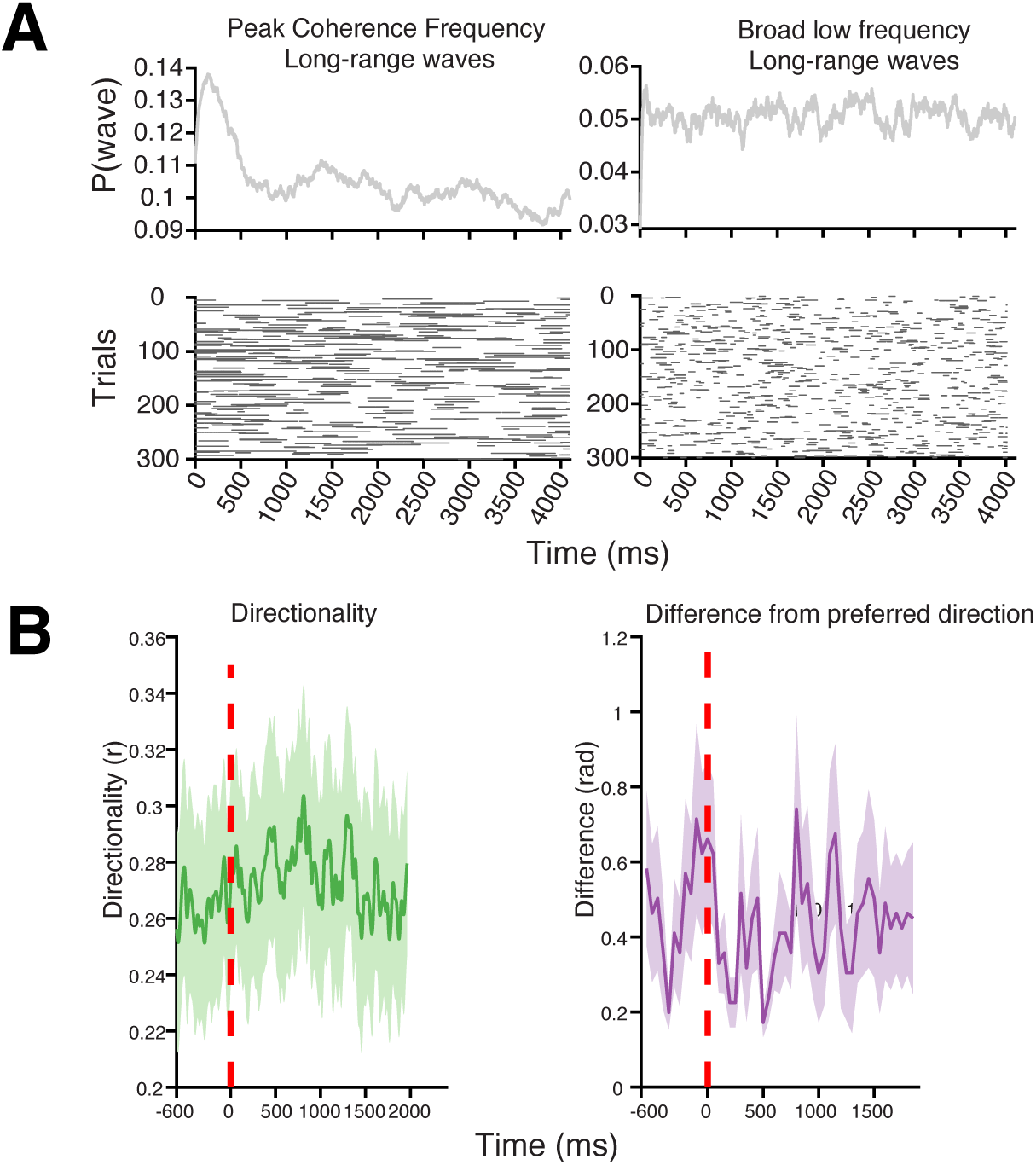
Filtering effects and artifact detection (*A*) Long-range wave times accumulated across trials and participants were used to compute the likelihood of observing a long-range wave (top panel). A random selection of 300 trials is shown in the raster on the bottom panel. The left plot shows a fixation offset-related (time = 0) increase in the probability of observing long-range frequency-specific waves. On the right is the corresponding plot for the 4 *−* 14 Hz long-range waves. Appropriate across-participant statistics are reported in Supplementary Information text. (*B*) The average directionality across participants of short-range waves in a 4 *−* 14 Hz broad low frequency band on the anterior temporal lobe and the average difference across participants between wave directions and the preferred direction (see Methods).

**Figure S8.**
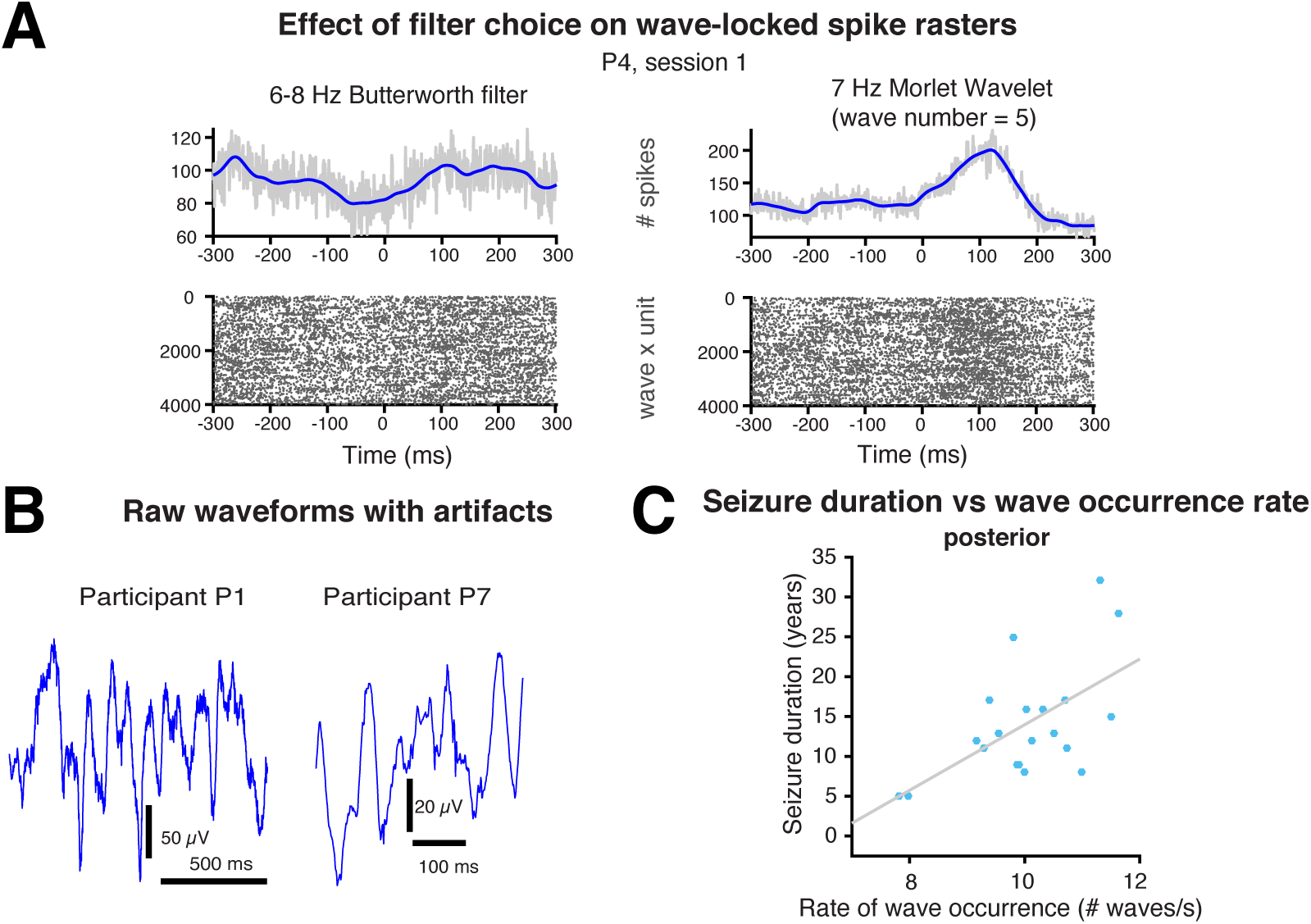
Filtering effects and artifact detection (*A*) Micro-scale wave-locked spike rasters for participant P4 and experimental session 1 using different filters, a spectrally precise but temporally imprecise second order 6 *−* 8 Hz Butterworth filter in one case and a temporally precise but spectrally relatively imprecise 7 Hz Morlet wavelet (wave number = 5) in the other. (*B*) Example artifactual epochs identified by the trial-level artifact rejection algorithm based on Staresina et al. (2015). (*C*) The relationship between seizure duration (in years) and rate of wave occurrence on the posterior 4 *×* 4 subgrid of electrodes.

**Table 1.**
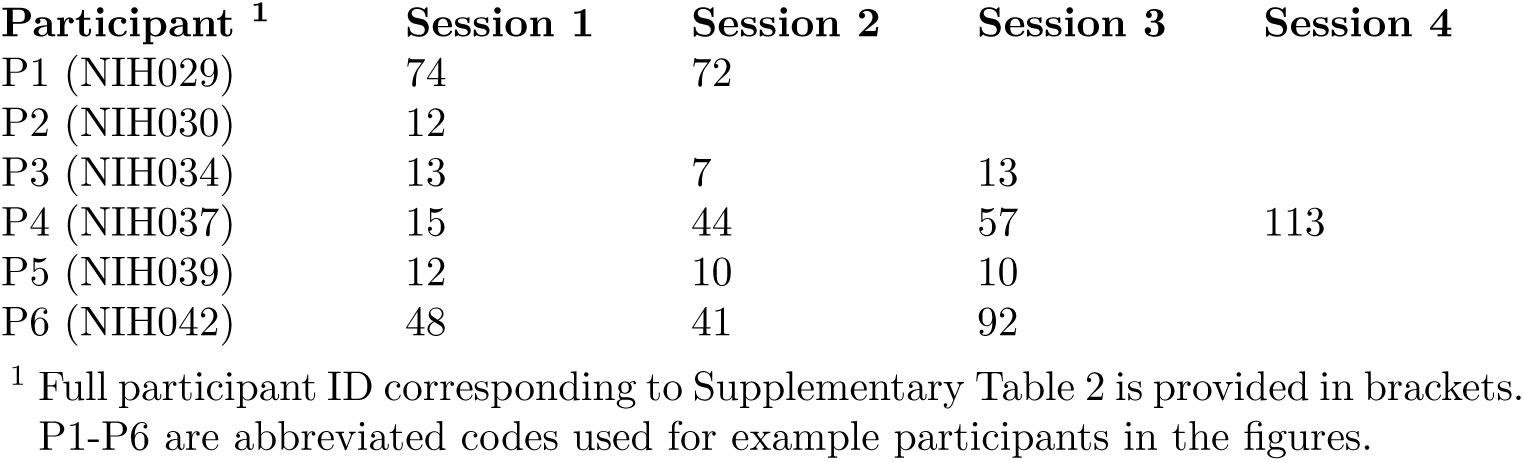
Number of well isolated-units detected in each experimental session and participant.

**Table 2.** Information about participants and epilepsy.

| **Participant ID** | **Sex** | **Procedure<sup>1</sup>** | **Age** | **Onset age** | **Seizure Frequency** | **Medication<sup>2</sup>** |
| --- | --- | --- | --- | --- | --- | --- |
| NIH007 | M | TOP | 25 | 13 | weekly | VPA 1000 BID |
| NIH011 | F | ATL | 43 | 38 | weekly | LTG 150 BID, ZNS 150 BID |
| NIH012 | F | ATL+TOP | 54 | 22 | weekly | CBZ 800 BID, LEV 750/1500 |
| NIH015 | M | NR | 25 | 17 | daily | LTG 450 BID, LCS 400 BID |
| NIH016 | F | TOP | 33 | 18 | weekly | LEV 1500 BID, TPM 75 BID |
| NIH018 | F | SAH | 30 | 14 | monthly | LTG 200 TID, TPM 50 QHS |
| NIH022 | F | ATL | 46 | 38 | monthly | LEV 2000 BID, LCS 200 BID |
| NIH024 | F | ATL | 29 | 17 | daily | LCS 150/150/300, CLB 20 BID |
| NIH026 | F | ATL+TOP | 34 | 6 | weekly | ESL 1200 BID, ZNS 200 BID |
| NIH029 | M | NR | 28 | 15 | monthly | LEV 2000 BID, LTG 100 BID |
| NIH030 | M | ATL | 39 | 14 | daily | CBZ 600 BID, LCS 200 BID |
| NIH032 | F | ATL | 35 | 19 | unk | LCS 200/300, LTG 75 BID |
| NIH034 | M | SAH | 51 | 40 | daily | LEV 500 BID, LCS 200/400, ESL 400 |
| NIH035 | F | TOP | 43 | 32 | weekly | LEV 2000 BID, LCS 200 BID, ZNS 200 BID |
| NIH036 | M | ATL+nodules | 33 | 24 | weekly | LEV 1500 BID, LCS 150 BID |
| NIH037 | F | ATL | 58 | 41 | weekly | LTG 100 BID, CBZ 400 BID |
| NIH039 | F | ATL | 28 | 19 | weekly | LEV 1500 BID, LCS 200 BID, TPM 25 BID |
| NIH042 | M | TOP | 20 | 7 | monthly | LEV 2000 BID, TPM 200 BID |
| NIH045 | M | TOP | 27 | 22 | daily | LCS 300 BID, CBZ 400 BID |
| NIH047 | M | ATL+nodules | 32 | 15 | weekly | LEV 1500 BID, PHT 200 BID |

<sup>1</sup> TOP = topectomy, ATL = anterior temporal lobectomy, SAH = selective amygdalohippocampectomy, NR = no resection;
<sup>2</sup> VPA = valproic acid, LTG = lamotrigine, ZNS = zonisamide, CBZ = carbamazepine, LEV = levetiracetam, LCS = lacosamide, TPM = topiramate, CLB = clobazam, ESL = eslicarbazepine, PHT = phenytoin.

